# Bayesian segmentation of spatially resolved transcriptomics data

**DOI:** 10.1101/2020.10.05.326777

**Authors:** Viktor Petukhov, Ruslan A. Soldatov, Konstantin Khodosevich, Peter V. Kharchenko

## Abstract

Spatial transcriptomics is an emerging stack of technologies, which adds spatial dimension to conventional single-cell RNA-sequencing. New protocols, based on *in situ* sequencing or multiplexed RNA fluorescent *in situ* hybridization register positions of single molecules in fixed tissue slices. Analysis of such data at the level of individual cells, however, requires accurate identification of cell boundaries. While many existing methods are able to approximate cell center positions using nuclei stains, current protocols do not report robust signal on the cell membranes, making accurate cell segmentation a key barrier for downstream analysis and interpretation of the data. To address this challenge, we developed a tool for Bayesian Segmentation of Spatial Transcriptomics Data (*Baysor*), which optimizes segmentation considering the likelihood of transcriptional composition, size and shape of the cell. The Bayesian approach can take into account nuclear or cytoplasm staining, however can also perform segmentation based on the detected transcripts alone. We show that Baysor segmentation can in some cases nearly double the number of the identified cells, while reducing contamination. Importantly, we demonstrate that Baysor performs well on data acquired using five different spatially-resolved protocols, making it a useful general tool for analysis of high-resolution spatial data.

---

During the last decade, single-cell transcriptomic technologies gained great popularity, with singlecell RNA-sequencing (scRNA-seq) has become the preferred approach for characterizing the state of complex tissues^1–4^. These techniques are being gradually augmented by the spatially-resolved transcriptomics measurements, based on *in situ* sequencing^5,6^, multiplexed single-molecule fluorescent *in situ* hybridization (sm-FISH)^7–9^, or spatially-barcoded hybridization^10,11^. The ability to examine the physical positions of different transcripts and cells at genomic scales has potential to bridge the molecular view of the cell with morphology, electrophysiology and other cellular phenotypes^12^. It can expose the impact of physical and biochemical interactions between cells, and reveal how such processes influence tissue organization during development^13^ and disease^14^. These protocols may eventually supplant scRNA-seq, as they also offer technical advantages, such as the ability to bypass capricious tissue disassociation steps needed for scRNA-seq. At present, however, most such assays are limited in the number of genes they can detect (30-300 genes), as well as the number of molecules that can be detected per cell (50-500)^8,15^. Nevertheless, there has been steady progress on the optimization of these protocols, with some increasing the number of detectable genes to thousands^7,16^. Increasing scales and spatial resolution are already enabling unbiased characterization of tissue organization^9^ and subcellular organization of cells^7,17^.

The transcriptional data acquired by the *in situ* sequencing or smFISH protocols can be generally summarized as a collection of detected molecules, each corresponding to a particular gene or transcript, along with the coordinates of that molecule within the field of view. While in principle such data can yield cellular or even sub-cellular resolution, the effective spatial resolution depends on the ability to distinguish features in the downstream analysis. Very sparse measurements, for instance, may only allow for interpretation of regional differences, such as segmentation of cortical layers. Achieving cellular resolution, however, even with high-density measurements, requires accurate cell segmentation. Most current groups have relied on the auxiliary nuclei staining (*e.g*. DAPI) to identify putative cell centers^7,9,16^. Unfortunately, even such one-channel segmentation is challenging, commonly requiring manual tuning and corrections^18^, including compensation for physical misalignment of molecular and auxiliary stains. The nuclei positions also do not inform on the extent of the cell body. Some efforts have used additional poly-A staining to extend the initial nuclei positions^9,16^. Similarly, pciSeq algorithm^15^ relies on the initial nuclei segmentation as a seed to extend the boundaries of the cell based on a Poisson model of gene expression. Alternatively, the spatial measurements can be analyzed without explicit cell segmentation (segmentation-free). Such approaches can characterize cell type composition of the tissue or identify distinct regions, but cannot be easily extended to many other kinds of downstream analyses^13,19^.

In this manuscript we start by discussing the applications and limitations of the segmentation-free approach. We suggest a new, simple method for segmentation-free analysis, which does not require extensive hyperparameter tuning. We then describe a general framework, based on Markov Random Fields, that can be used to solve a variety of molecule labeling problems. In particular, we follow this strategy to implement solutions for (i) separation of background noise, (ii) *de novo* inference of cell populations without cell segmentation, and (iii) cell type annotation of molecules based on the annotated scRNA-seq data. Finally, we introduce a method that performs cell segmentation based on the observed molecules, and optional microscopy staining data. All of the algorithms are implemented in an open-sourced command-line tool and a corresponding Julia package called *Baysor*. We show that Baysor can segment data from most published protocols with molecular resolution, yielding better segmentation accuracy, increasing the number of detected cells, as well as the number of molecules associated with each cell.

## Results

### Analysis of local expression patterns without segmentation

As illustrated by the scRNA-seq studies, different cell types and many phenotypic states can be readily distinguished based on the transcriptional composition of a cell. In spatial measurements, the cells of a distinct type will give rise to small molecular neighborhoods with stereotypical transcriptional composition. This patch-like structure of the spatial transcriptomics data can be used to interpret it without performing explicit cell segmentation^19^. To perform such neighborhood composition analysis, we generated a neighbourhood composition vector (NCV) for each molecule by taking its *k* spatially nearest neighbors and estimating the relative frequency of different genes among the neighboring molecules (Fig. 1a). These expression vectors can be treated as “pseudo-cells” and analyzed using existing methods developed for scRNA-seq, including clustering, cell type annotation and embedding (Fig. 1b,c). The NCVs can also be used to effectively visualize local transcriptional composition. To do so, we embed the NCVs in 3D color space. Under such color encoding, where the neighborhoods of similar transcriptional composition are represented by similar colors, different types of cells as well as their boundaries become visually apparent (Fig. 1).

**Fig. 1.**
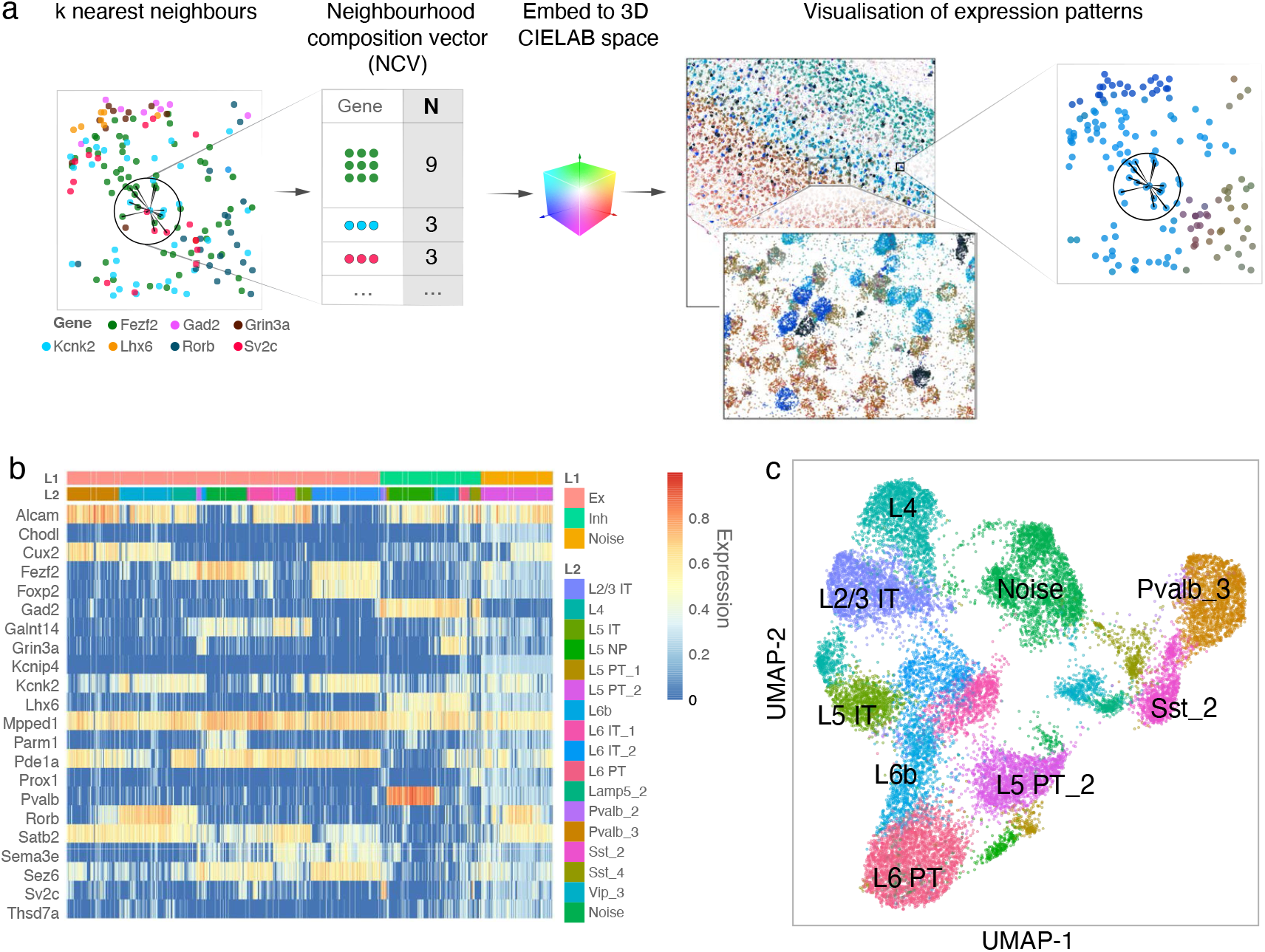
Segmentation-free analysis of spatial data using Neighbourhood Composition Vectors (NCVs). **a**. NCVs are estimated by taking *k* spatially-nearest neighbours for each molecule, and tabulating the number of neighborhood molecules belonging to each gene. (left) The molecules measured in the 2D space are shown as dots, colored by the gene identity. Based on the similarity of these vector profiles, NCVs can be clustered or embedded into lower-dimensional space (b,c). (center, right) A 3D embedding can be translated into a color encoding, so that similar colors correspond to similar neighbourhood compositions. Such color encoding allows for effective visualization of individual cells as well as the overall tissue organization. A part of the Allen smFISH dataset is shown as an example. **b**. The heatmap shows the expression patterns for 20000 NCVs, uniformly sampled across the physical space with rows corresponding to genes and columns corresponding to NCVs. The color scale shows log_10_ of total-count normalised expression, additionally normalised by the maximum for each gene. The L1 and L2 column headers show the marker-based annotations for the corresponding NCVs. **c**. NCVs can be analyzed using existing scRNA-seq pipelines, to generate clustering, cell type annotation and embeddings. An 2D UMAP embedding is shown, labeled and colored according to the published annotation of the corresponding cells.

Unlike scRNA-seq datasets, spatial transcriptomics data can be analyzed at different scales. In the NCV analysis, the spatial scale is determined by the neighborhood size parameter *k*, relative to the average number of molecules measured per cell *m*. Small neighborhoods, with characteristic size smaller than the scale of the cell (*k ≪ m*), can provide information about intra-cellular organization, driven for example, by the nucleus or other organelles. Most of the protocols published to date, however, lack the resolution necessary to effectively distinguish such subcellular features. The notable exceptions are the Seq-FISH+^7^ protocol and a high-resolution variant of MERFISH^9,20^. Large neighborhoods, with characteristic size greater than a cell (*k ≫ m*), can reveal microanatomical tissue and even organ-level organization, such as the layer structure of the brain cortex (Supplementary Figs. 1 to 3).

### General approach for statistical labeling of spatial data

A number of analyses in spatial transcriptomics can be formulated as label-assignment problems. Cell segmentation, for instance, assigns cell labels to the observed molecules. Similarly, separation of intercellular background is a problem of labeling molecules as “signal” *vs*. “background”. The distinguishing characteristics of these problems is that the labels tend to show strong spatial clustering: two nearby molecules, for instance, are likely to belong to the same cell and therefore share a common label. Mathematically, this spatial clustering tendency can be captured using Markov Random Field (MRF) priors^21,22^. The labels themselves can be modeled as latent variables, and inferred from the observed data using an Expectation-Maximization (EM) algorithm.

Different labeling problems can then be solved by choosing the appropriate label probability model and the observable data (Supplementary Fig. 4a). For instance, by using gene identities of the molecules as observables, and multinomial distributions to model the transcriptional composition associated with different labels, this MRF-based approach results in a meaningful clustering of molecular neighborhoods (Fig. 2a,b). One can additionally use expression profiles of different cell types obtained from scRNA-seq data as a prior for the multinomial distributions of different labels. This enables the approach to efficiently transfer the cell annotations from scRNA-seq to the measured molecules without performing cell segmentation (Supplementary Fig. 5). The MRF-based inference can be notably faster than traditional clustering (Supplementary Table 1), however, both performance and robustness of such annotation transfer become poor when the number of cell types increases beyond 10-20. Another example of the labeling problem is distinguishing background molecules from cell bodies. In this setting, one can assume that the cells form dense regions, while the background noise molecules appear in sparse regions. Taking the distance to the *k*-th nearest neighbor as a measure of sparsity (observed data), we used the same EM algorithm to segment the background (Fig. 2c-d and Supplementary Fig. 4a). Overall, MRF provides a general recipe for solving a variety of spatial labeling problems, though each problem requires a custom formulation of the EM algorithm.

**Fig. 2.**
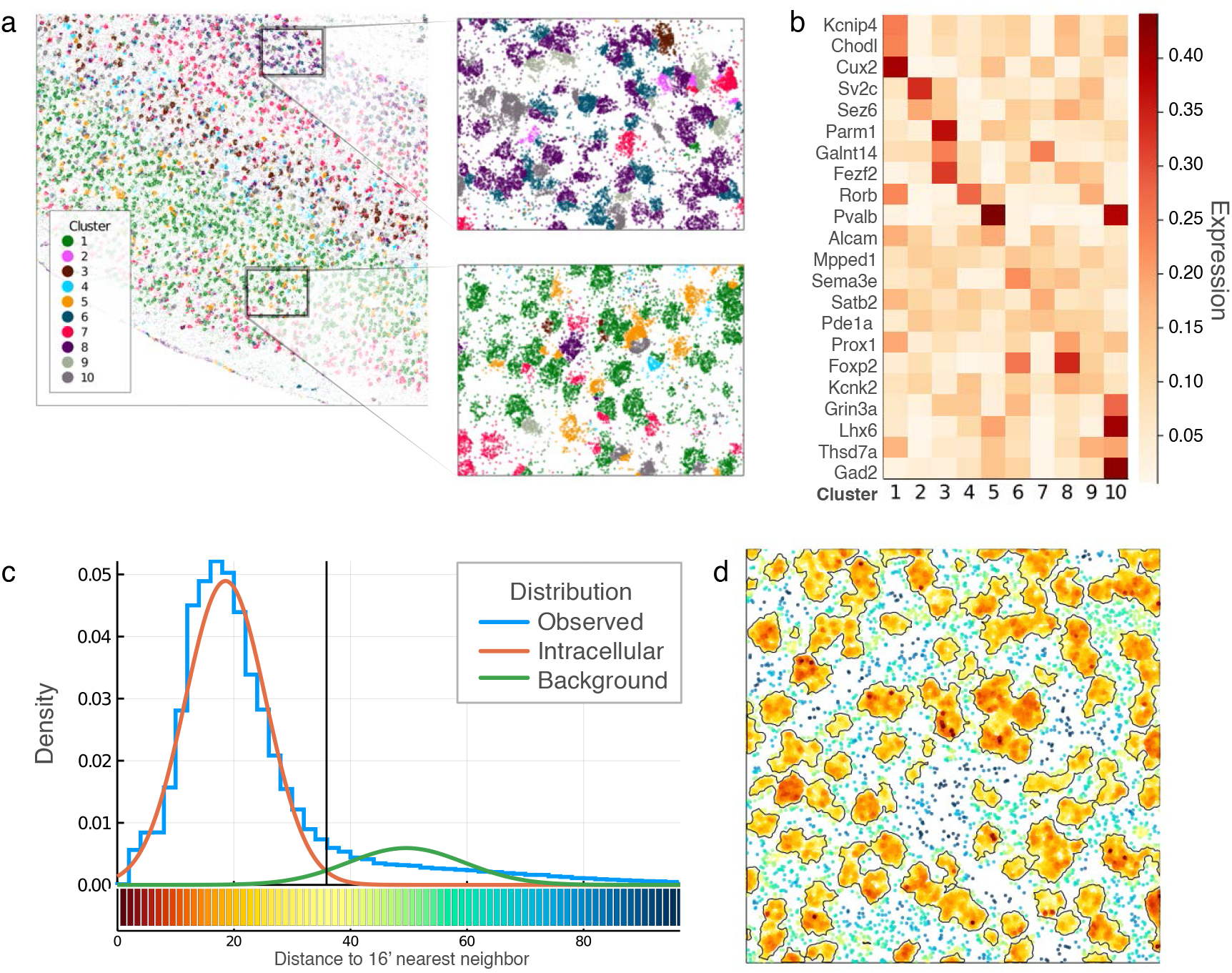
Application of Markov Random Field (MRF) framework for segmentation-free cell type inference and background filtration. **a,b**. Individual molecules were clustered into major cell types by modelling the tissue as a mixture of multinomial distributions with MRF prior. Cluster labels per molecule are shown in (a), with expression vector for each of the clusters shown in (b). **c**. The MRF approach is used to separate background from intracellular signal. For each molecule, the algorithm estimates the distance to its *k*-th nearest neighbour and models the distances as a mixture of two Normal distributions. The distribution of physical distances to 16-th nearest neighbour (x-axis) is shown for the molecules in the Allen-smFISH dataset as a histogram (blue). Fitted intracellular and background distributions are shown in red and green, respectively. The vertical black line shows the optimal separation point. **d**. Molecules from a subset of the Allen-smFISH dataset are shown as dots, coloured by their distance to the 16-th nearest neighbour, with the colorkey shown on the bottom of (c). The black contours mark regions above 50% probability of being intracellular.

### Cell segmentation across various protocols

Despite the relative ease and effectiveness of the NCV approach described above, many of the downstream analyses and interpretations of the spatially-resolved data depend on the ability to resolve individual cells. These include analysis of context-dependent cell expression states, physical interactions and spatial dependencies between cell types, cell migration and formation of tissue architecture. We, therefore, set out to develop a cell segmentation method that can take into account different facets of data that are informative of cell boundaries. The increased spatial density of molecules within the cell somas is one such facet. The transcriptional composition of local molecular neighborhoods is another. Further evidence can be gained from stainings for nuclei (*e.g*. DAPI), cell bodies (*e.g*. poly-A primers), or cellular membranes. To optimize cell segmentation based on multiple evidence sources, we have developed an algorithm, called Baysor, which builds on the ideas of the MRF segmentation outlined above. The method can be used to analyze data from various experimental protocols (Fig. 3), and can perform cell segmentation using molecular positions alone, or by incorporating additional information. The approach models each cell as a distribution, combining spatial positions and gene identity of each molecule. Thus, the whole dataset is considered as a mixture of such cell-specific distributions. Baysor then uses Bayesian Mixture Models to separate the mixture. The optimization relies on MRF prior to ensure spatial separability of the cells and to encode additional information about the spatial relations of molecules (Supplementary Fig. 4b).

**Fig. 3.**
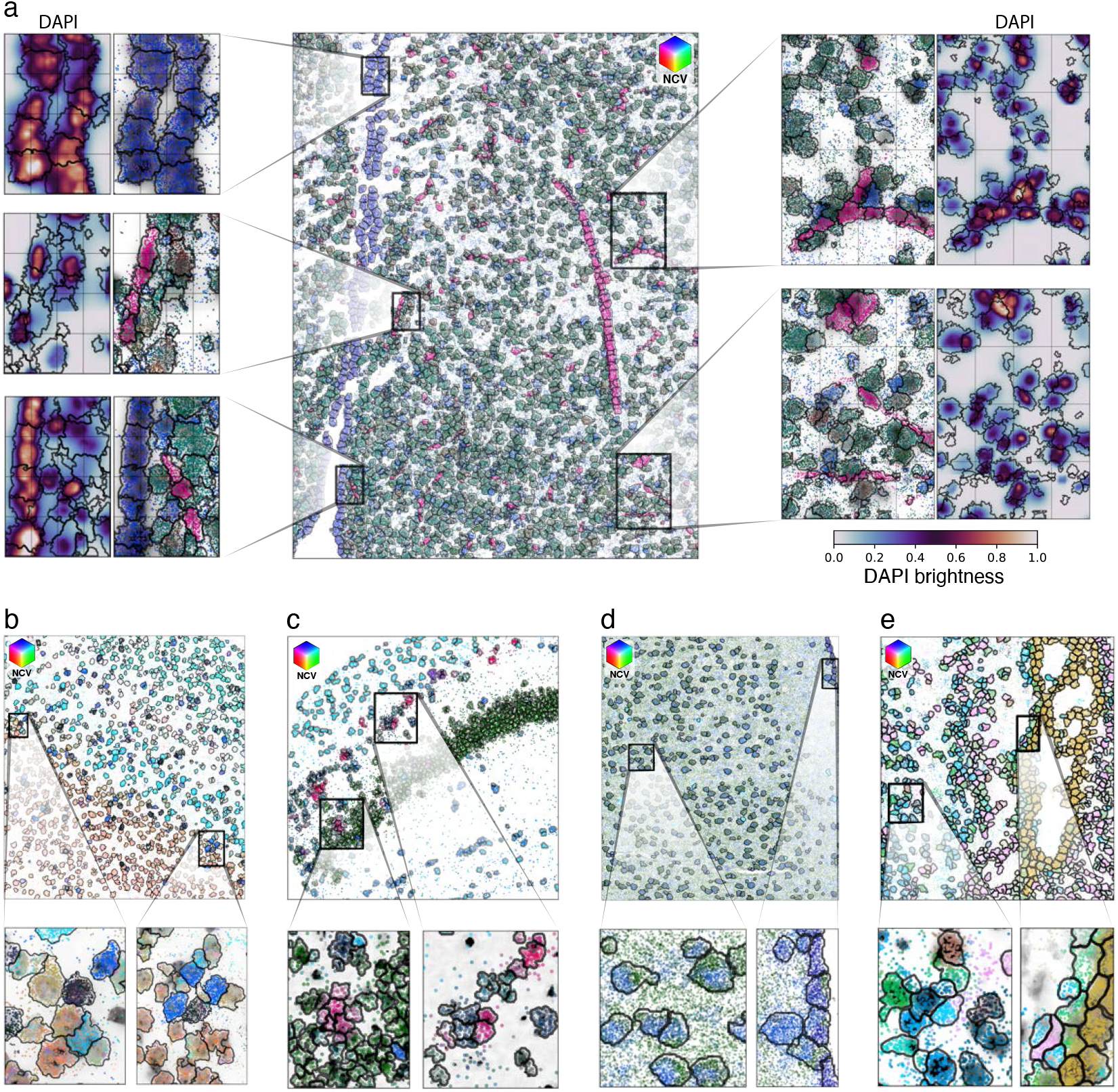
Examples of Baysor cell segmentation over the published protocols. **a**, Baysor segmentation is shown for a part of the MERFISH Mouse Hypothalamus^9^ dataset. The central panel shows positions of the measured molecules, colored by their neighbourhood gene composition (see Fig. 1a). The inferred boundaries of the segmented cells are shown as black contours. Zoom-in views are shown immediately to the left and right of the central plot. The outer plots show DAPI signal within these regions. Additionally, the DAPI signal is also shown as grayscale background within the zoom-in molecule plots. **b-e**, Additional examples of Baysor segmentation are shown for the Allen sm-FISH Mouse VISp (**b**), ISS Hippocampus^15^ (**c**), STARmap Mouse VISp 1020^16^ (**d**), and osm-FISH somatosensory cortex^8^ (**e**) datasets.

Existing cell segmentation methods rely on nuclear (DAPI) or cytoplasmic (poly-A) staining^8,9,15^, segmenting the images with watershed or other algorithms to obtain cell labels^18,23^. While Baysor can perform segmentation using only the information on the measured molecules (Fig. 2), the auxiliary stains can provide valuable information in cases where the molecular signal is sparse or not informative about cell boundaries (see Discussion). Baysor can take advantage of such information by using a pre-calculated segmentation as a probabilistic prior. Computational segmentation of nuclear and cytoplasmic stains, however, remains a challenge in itself^18,19^, and the pre-calculated segmentations will typically contain many errors (Fig. 4d). To account for this, Baysor defines a “prior segmentation confidence” parameter which determines the weight of the prior. Setting this parameter to 0 will cause Baysor to ignore the prior, while a maximum value of 1 will restrict Baysor from changing segmentation of the molecules assigned to cells, leaving it to deal only with non-assigned molecules (Supplementary Fig. 6). Prior segmentation is also taken into account when determining the background to penalize removal of the molecules assigned to cells in the prior segmentation (see Methods). In addition to segmentation priors, Baysor can also incorporate information about background assignment probabilities per molecule. Finally, Baysor can use information about molecule clustering to penalize assignment of molecules from different clusters to the same cell.

**Fig. 4.**
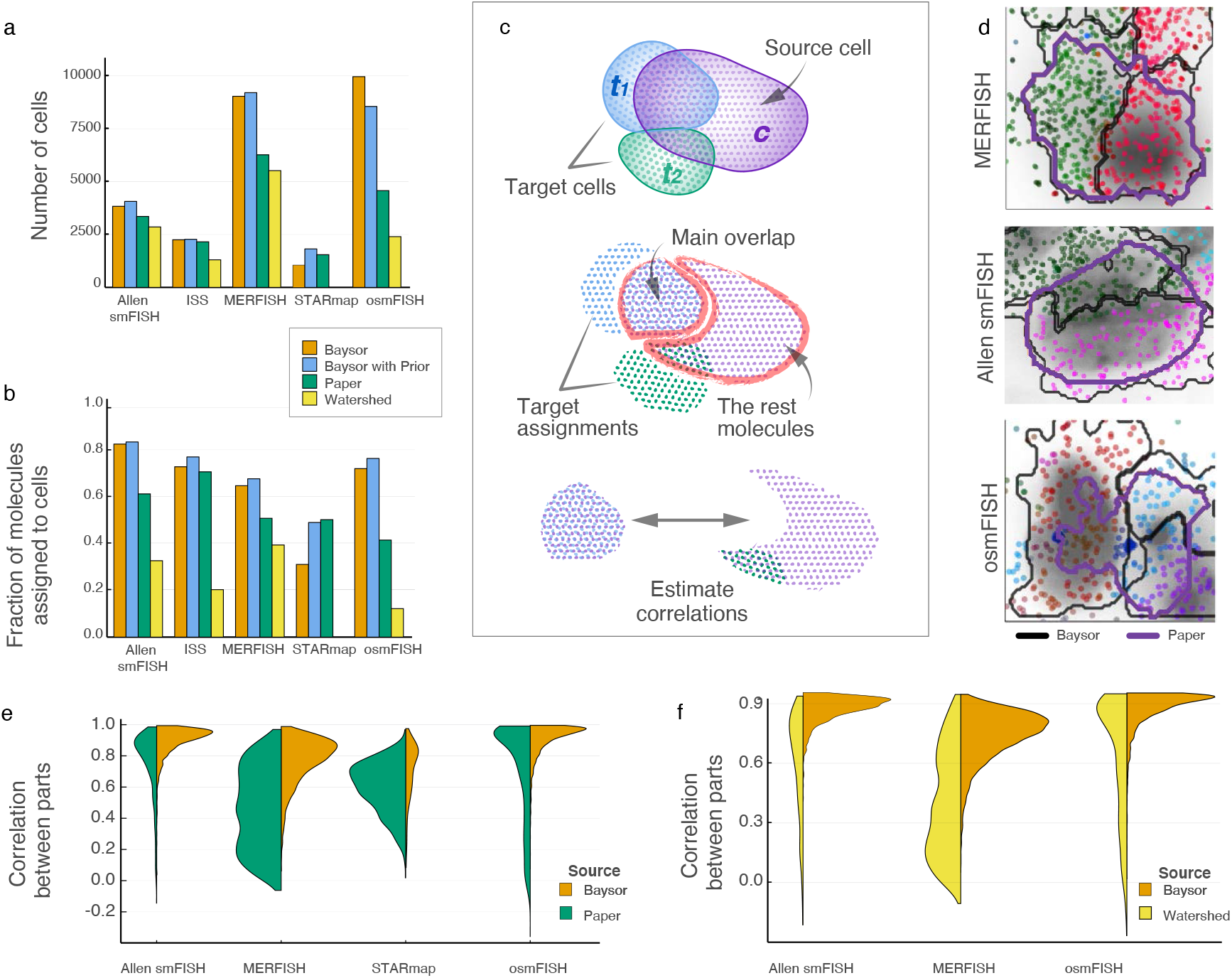
Comparison of the Baysor segmentation to the published results and Watershed DAPI segmentation. **a,b**, Number of cells (**a**), and fraction of the molecules assigned to cells (**b**) by different segmentation methods (color) are shown for different datasets (x-axis). **c**, Schematics for evaluating the differences between two segmentations based on gene composition of the results. Each cell from the source segmentation *c* is matched to cells from the target segmentation *t*. For the source cells, which overlap multiple target cells, the region with the largest (“main”) overlap is used. Correlation of gene expression of the main overlap region against the expression of the rest of the cell in the source segmentation is then estimated. **d**, Examples of the results where the published segmentation merged distinct cell types. The dots show the measured molecules, colored by NCVs with contours showing cell boundaries for Baysor (black) and reported in the original publications (purple). **e**, Comparison of Baysor results to the published segmentations, using the correlation benchmark (**c**). The violin plots show the distribution of overlap correlations with the rest of the cell (y-axis), for different datasets (x-axis). The right part of each violin plot was calculated using Baysor segmentation as a source and the published segmentation as a target, while the left parts were calculated by swapping the source and target segmentations. The width of the violin plots is proportional to the number of source cells that were matched to multiple target cells. The results show that Baysor segmentation (used as a target) can be used to split multiple cells from the published segmentations into poorly-correlated parts, while the reverse - using the published segmentation as a target - for the most part is not able to identify flaws in the Baysor segmentation. **f**, Analogous to **e**, the plot compares performance of Baysor segmentation with the segmentation obtained using Watershed algorithm.

To evaluate the performance of our approach, we compared Baysor results with segmentations provided in the original publications (“Paper”). Additionally, a common base-line segmentation was generated using a watershed segmentation of DAPI images (using ImageJ^23^, see Methods). Baysor was run in two configurations: a minimal configuration - using only the positions and gene identity of the detected molecules (“Baysor”); and using enhanced configuration where the originally published segmentations were used as a prior for Baysor segmentation (“Baysor with Prior”). We first examined various summary statistics for different segmentations. Both Baysor and the Paper segmentations have around the same number of molecules and area per cell (Supplementary Fig. 7), which suggests that neither of them performs over- or under-segmentation. In contrast, the Watershed segmentation has cells of smaller size, which can be explained by it capturing only the nuclei information and discarding cytoplasmic molecules. Compared to published (Paper) segmentations, Baysor reports a larger number of cells and a higher fraction of molecules recognized as a part of a cell (Fig. 4a,b) with the largest difference of 2 folds for the osmFISH data^8^. The Watershed underperforms other methods by these two criteria, as well. Additionally, since the Baysor algorithm is stochastic in nature, we showed that the segmentations generated based on different seeds of the random number generator showed highly similar (Supplementary Fig. 8). Additionally, we profiled time and memory usage of the Baysor run with the longest run taking 51 minutes for the MERFISH dataset with 3.7M molecules and the largest memory usage of 40.4 GB for the STARmap dataset with 1020 genes (Supplementary Table 2).

We currently lack experimental methods to establish ground truth on cell segmentation, so it is not possible to estimate a global quality metric which would show to what extent the results differ from an ideal segmentation. Instead, we examined the differences between segmentations and evaluated which algorithm performs better in the cases where the segmentations disagree. Specifically, in comparing any two segmentation results we identified all cases where a cell from one segmentation matched multiple cells from the other segmentation. For each of these cases we picked the largest part of the cell from the first segmentation that matched to a single cell from the second segmentation. We then estimated the correlation of expression profiles between this matching part and the rest of the cell (Fig. 4c). If the first segmentation was correct, then the matching part should show similar transcriptional composition to the rest of the cell in the first segmentation, and the resulting correlation measure will be high. In contrast, if the second segmentation was correct, the expression correlation will be low (Fig. 4d). Assessment of such expression correlation between parts of the cell requires a relatively high number of molecules per cell, so we were not able to apply this benchmark to the dataset generated using ISS protocol^15^ (Supplementary Fig. 7a). On all other protocols, the overlapping regions showed on average higher expression correlation with the corresponding Baysor assignments than with the alternative segmentations (Fig. 4e,f and Supplementary Fig. 9), indicating higher accuracy of Baysor segmentation results.

We further investigated the two datasets where the differences between Baysor and published segmentations were most notable: osmFISH^8^ (Fig. 5) and MERFISH^9^ (Supplementary Fig. 10) datasets. In both cases, the segmentation differences preferentially impacted certain cell types. In the case of osmFISH, the published segmentation omitted most of the cells of non-neuronal subtypes: only 8% of Vascular and Astrocytic cells detected by Baysor are present in the original segmentation (Fig. 5d). The disagreements on the MERFISH dataset were less biased, with the largest difference observed for the Endothelial cells, with the published segmentation reporting 51% fewer cells (Supplementary Fig. 10c). There were two subtypes where Baysor distinguished fewer cells: 10% less for Ependymal, and 15% for Microglia. The difference in Microglia, however, is likely caused by the set of Inhibitory neurons that express contradictory markers and were mis-annotated as Microglia (Supplementary Fig. 10j).

**Fig. 5.**
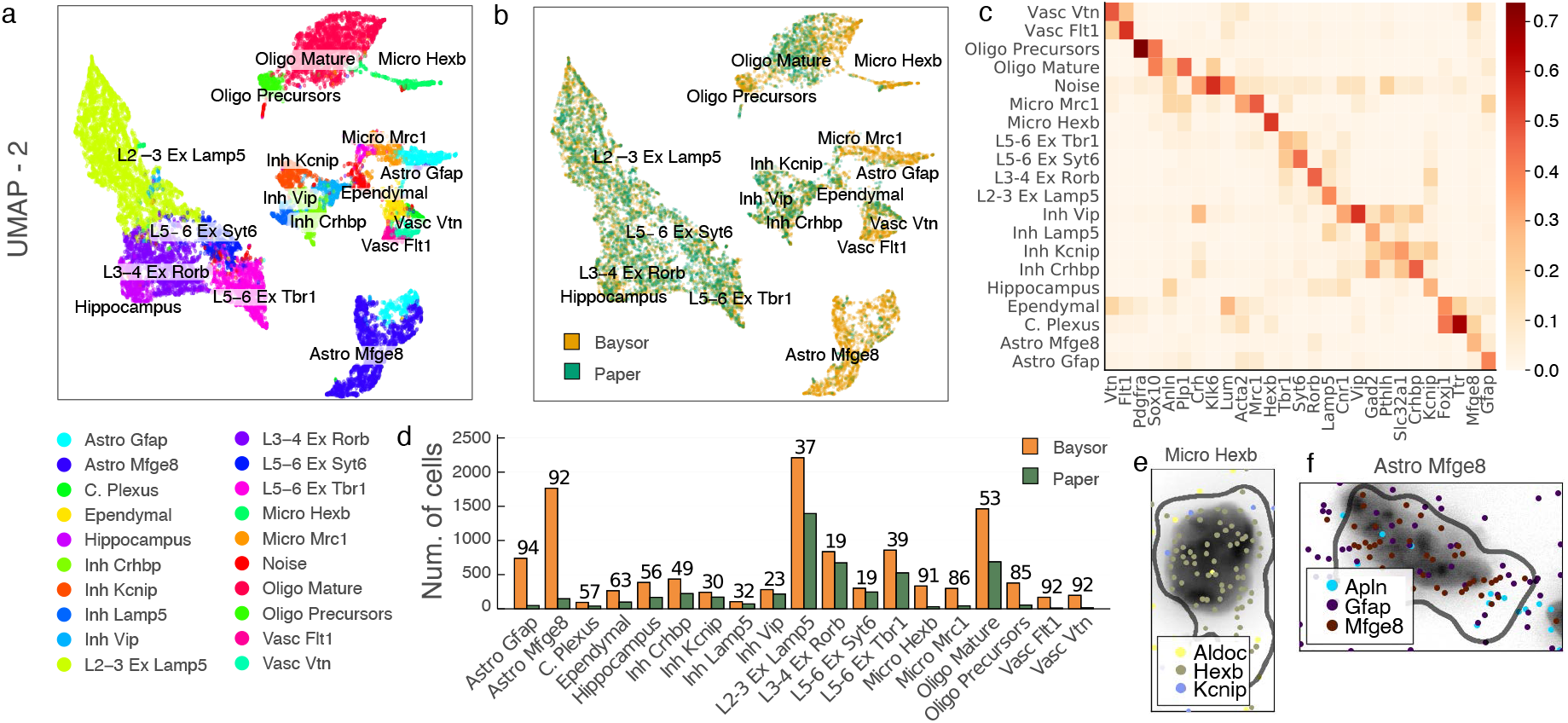
Examples of the segmentation differences on the osmFISH data. **a**, A joint UMAP embedding of the cells generated by both Baysor and the published segmentations, labeling the annotated cell types with color. **b**, The same embedding, colored by the segmentation method. **c**, A heatmap showing expression patterns of marker genes (columns) for each of the cell types (rows). The colours show expression levels, normalised by gene. **d**, The barplots showing the number of cells per cell type for the Baysor (brown) and the published (green) segmentations, with the numbers on the top of the bars showing excess percentage for the Baysor segmentation. **e-f**, Examples of Micro Hexb (**e**) and Astro Mfge8 (**f**) cells, which were not segmented in the published segmentation, but were distinguished using Baysor. The dots correspond to molecules, colored by gene (only five the most abundant genes are shown). The grayscale background represents DAPI signal, and the black line shows the cell boundary determined by Baysor.

### Outstanding challenges

The Baysor model described above performed well on most existing protocols, however some edge cases are not resolvable within the current model. These include working with ultra-high resolution data, capturing 3D structure of the data, and segmenting sparse homogeneous regions. An example of the ultra-high resolution data is the NIH3T3 fibroblast dataset from the Seq-FISH+ protocol^7^. It captures 10,000 different genes with 35,622 molecules and 6,700 unique genes per cell on average. Such resolution reveals prominent sub-cellular structure, resulting in heterogeneous gene composition within different parts of the cell body (Fig. 6a). Furthermore, such data also highlights complex morphology of the cells (Fig. 6b). These features go beyond the two critical assumptions of the current Baysor model: that the composition of the cell body is homogeneous, and that the cell shape can be reasonably well approximated using a bivariate normal prior. The homogeneity assumption was also violated in the dataset generated using STARmap protocol^16^, however the reasons for that are likely technical: in the STARmap data, the molecules belonging to the same gene were spatially clustered, with the size of such mono-genic clusters reaching dozens of molecules (Fig. 6d,e and Supplementary Fig. 11). While Baysor was still able to perform a reasonable segmentation of this data (Fig. 4a,b,e), such local clustering of genes affected the convergence of the algorithm.

**Fig. 6.**
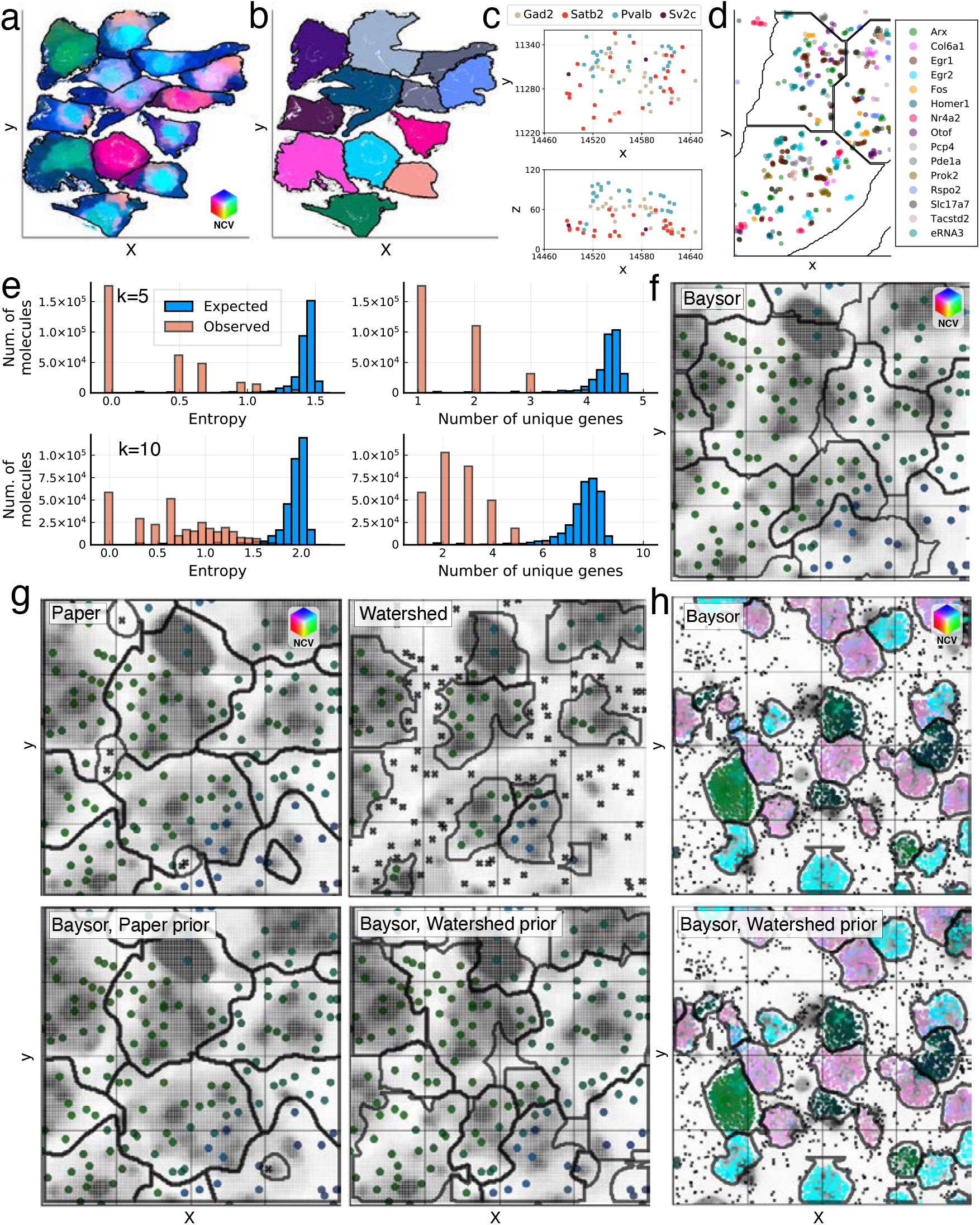
Outstanding challenges. **a**, Seq-FISH+ Fibroblast^7^ data colored by NCVs with black contours showing the published segmentation borders. **b**, The same data, segmented by Baysor with colors showing cell assignment. **c**, Example of cells which are separable only in 3D in the Allen smFISH data. The two plots show 2D projections on the physical x-y and x-z axes correspondingly. Each point represents a molecule, coloured by its gene of origin. Gad2 and Pvalb are markers of inhibitory neurons, while Sv2c with Satb2 are markers of excitatory neurons. These markers are mutually exclusive, and there should be no cell that expresses all four of these markers. **d**, Example of a cell from the STARmap VISp 160 dataset^16^. The black lines show the published cell boundaries. The plot shows colouring by gene for the 15 most expressed genes. **e**, Local grouping tendency of the transcripts on STARmap data, is illustrated through distributions of neighbor gene entropy. (top) For each molecule, *k* = 5 nearest neighbours were estimated. The entropy (left) of their gene count vector, and the number of unique genes in the neighbourhood (right) and are shown on the “Observed” histogram. The “Expected” histogram shows the distributions expected under gene randomization. (bottom) Analogous plots for k = 10 neighborhoods. **f**, Example of a homogeneous CA1 regionin the ISS data^15^. The plot shows Baysor segmentation, and it can be seen that cell boundaries do not match the grayscale DAPI signal from the nuclei. Each dot represents a molecule, colored by NCVs, with black contours showing cell boundaries. **g**, Different segmentations shown for the same region as **f** with the segmentation type specified in the topleft corner. Black crosses on the plot show molecules, assigned to background. The bottom row shows that after using DAPI segmentation (Watershed) prior, Baysor segmentation shows better correspondence to the nuclei DAPI signal. **h**, Segmentation examples from the Allen smFISH dataset, showing Baysor segmentation without (top) and with (bottom) DAPI-based Watershed prior. Here, using Watershed prior does not change results visibly, as transcriptomic signal matches to DAPI.

Another potential limitation is the presence of the 3D structure in the data. The existing protocols can scan through the third dimension by generating sequential focal stacks (z stacks). The distance between the sequential stacks, however, can vary. In some datasets, the distance between stacks exceeds 10μm, effectively capturing different cell layers. Other datasets show much smaller distances, on the order of *1μm,* in which case the stacks can be pooled together reducing it to a 2D view. There are, however, some rare cases where the stacks capture real 3D structure of the cells (Fig. 6c). In such cases, segmentation needs to be performed in 3D. While the Baysor model can be extended to 3D, the current implementation is limited to 2D data.

The third challenge involves separation of sparse homogeneous regions. If a region is composed primarily of the same cell type, the segmentation would normally be driven by the local density clustering of the detected molecules (*e.g*. Fig. 6b). However, if the molecular signal is very sparse, such density patterns become challenging to detect. This situation can be observed in the ISS dataset of the CA1 brain region^15^. As there is little signal in the data, the resulting Baysor segmentation does not correspond to the DAPI staining (Fig. 6f). It is worth noting, however, that for the purposes of the downstream analyses, the uncertainty in the boundaries between cells of the same type is likely to be less consequential than other types of errors.

The challenging situations described above can be addressed by specifying a segmentation prior based on auxiliary stainings, and regulating its weight with the “prior segmentation confidence” parameter. For accurate segmentation of complex cell shapes, stainings that register the whole cell body are needed (*e.g*. poly-A or membrane staining). The choice of the staining segmentation method is also important, to ensure that the prior segmentation traces the complex cell shapes. When using such priors, the further the cell shapes are from simple ellipsoid approximations, the larger should be the value of the prior segmentation confidence parameter. Similarly, having whole body stainings with high prior confidence can help to overcome problems arising from heterogeneous subcellular structure. For instance, incorporating a segmentation prior for STARmap VISp 160 had a substantial effect on the segmentation (Supplementary Figs. 9, 12 and 13).

In contrast, segmentation of sparse datasets can be aided by the knowledge of cell centers alone, obtained from DAPI or similar stainings. Specification of the approximate cell size (*i.e*. global scale) is also useful in such cases. A lower prior confidence value will provide better results in such cases, since even a small value should be sufficient to resolve homogeneous regions. Using larger prior confidence value, on the other hand, can distort the segmentation of heterogeneous regions. Prior segmentations do not appear to help in overcoming 3D structure effects, as the current 2D model can not properly account for such data.

## Discussion

Realizing the potential of spatially-resolved transcriptomics will require continued improvements on both the side of the protocols^12^, as well as analytical methods for processing such data. Here we focused on addressing an important pre-processing step of cell segmentation. Effective segmentation can increase the number of detected cells, and provide more informative profiles for each cell. The accuracy of the segmentation is also critical for a number of valuable downstream inferences. For instance, incorrectly drawn borders can create spurious correlation of expression state between adjacent cell types, resulting in false-positive inference of cell interactions. Alternatively, shifted borders may be interpreted as transient cell states, suggesting false transitions between cell types. To avoid such potential issues, we described an approach that uses transcriptional composition to optimize the placement of cell boundaries. Baysor can perform segmentation using only molecule placement data or in combination with evidence from auxiliary stains, and yields improved segmentation quality, increased number of cells and segmented molecules.

Not all of the downstream analyses require cell segmentation. For instance, region segmentation or tissue cell type composition may be inferred directly from molecular data^19^. We show that a relatively simple segmentation-free approach based on the composition of local neighborhoods (NCVs) can be used to assess the quality of the dataset, estimate the number and identity of the major cell types, and effectively visualize the organization of the tissue (Fig. 1a). This approach is fast and does not require parameter tuning, making it a convenient option for preliminary analysis. Furthermore, the NCVs can be fed directly into existing scRNA-seq analyses methods for integration with other datasets, annotation of cell types, etc. Many of these downstream problems, however, can be formulated as labeling problems, in which local continuity can be effectively captured using Markov Random Field (MRF) priors. Such MRF-based approach is at the core of Baysor cell segmentation method, but can also be used to solve other labeling problems, such as separation of signal from background molecules or clustering of NCVs. It can also be used for continuous labels, such as cellular response on an injury, for example modeling dependence on the distance from the site of the injury. The strategy can also be applied on the level of cells, for instance to identify larger tissue segments^22^.

Though Baysor algorithm performed well on most of the published protocols, a number of potential improvements could be introduced. For instance, the implementation can be extended to support segmentations in 3D space, without altering the logic of the underlying algorithm. A more complex problem would be improving modeling of cell shapes^24^, for instance by replacing current ellipsoid shape approximations by limiting the size and complexity of the cellular shapes. Further improvements could be gained by extending the hierarchical Bayesian model to introduce cell-type specific shape and composition characteristics. Finally, as we demonstrate, information from auxiliary stainings can be extremely valuable in resolving difficult cases. Improved stainings, such as those labeling cellular membranes, as well as improved methods for segmenting such images will likely be key for improving the overall segmentation results. However, even with the common DAPI images, manual processing is commonly required to perform initial nuclei segmentation. As Baysor can take advantage of uncertain prior predictions, a probabilistic auxiliary image segmentation method that could incorporate nuclei, cytoplasm and membrane stainings to predict a cell center and boundary probability maps would provide a significant advantage. We hope that the Baysor implementation and the MRF-based computational approach will further facilitate the development and applications of high-resolution spatial transcriptomics methods.

## Acknowledgements

We thank Bosiljka Tasic and Brian Long for sharing the non-published Allen sm-FISH data and aiding in its interpretation, as well as the SpaceTx consortium for facilitating the collaborations. We would like to thank Yuri Boykov (U. of Waterloo) for the initial discussions and advice on an alternative segmentation approach based on graph cuts. We are also grateful to a number of colleagues who advised us on the published protocols: Jeff Moffit (MERFISH), Nico Pierson and Long Cai (Seq-FISH+), Simone Codeluppi, Lars Borm and Sten Linnarsson (osm-FISH), Xiaoyan Qian, Markus Hilscher and Mats Nilsson (ISS), as well as to Jeremy Miller for his input on segmentation benchmarks. We would like to express our gratitude to Dmitry Molchanov and Dmitry Vetrov (HSE, Moscow) for their input on the algorithm.

## Competing interests

P.V.K serves on the Scientific Advisory Board to Celsius Therapeutics, Inc. Other authors declare no conflict of interest.

## Methods

### Neighborhood Composition Vector Analysis

To analyze the spatial expression patterns without cell segmentation, we use neighbourhood composition vectors (NCVs) as a unit of analysis. NCVs are constructed by identifying *K* spatially nearest neighbours (NNs) for each molecule, and then characterizing its composition. That is, estimating the frequency of occurrences of different genes among the neighbours:

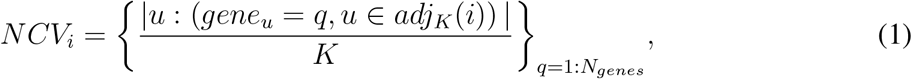

where *N_genes_* is the total number of measured genes, *gene_u_* is the gene that produced the molecule *u, K* is the number of NNs and *adj_K_* (i) are the indices of these NNs for the molecule *i*. To estimate the NNs, a k-d tree structure implemented in the NearestNeighbors.jl julia package was used. 2D Euclidean distance was used as a distance metric. As implemented in the Baysor package, the default value of *K* was set to the expected minimal number of molecules per cell (a user-modifiable parameter) or the total number of detectable genes divided by 10, whatever is larger.

To perform single-cell RNA-sequencing (scRNA-seq) analysis of NCVs we used Pagoda2 (https://github.com/kharchenkolab/pagoda2) R package to calculate UMAP embedding^25^ and CellAnnotatoR package (https://github.com/khodosevichlab/CellAnnotatoR) to annotate the cell types.

To visualize the local gene composition, embedding of the NCVs into three dimensions was performed, first by reducing the dimensions using Principal Component Analysis (PCA) to the top 15 principal components, and then embedding the data into 3D space using UMAP with the default parameters *min_dist* = 0.1 and *spread* = 2.0. As fitting UMAP embedding for many NCVs is computationally intensive, to optimize performance we first fit the UMAP embedding on a subset of NCVs and then applied the resulting transformation to all NCVs. First, PCA was estimated on the whole dataset. Then, 10000 molecules were selected uniformly across the principal components. After which UMAP was fitted only on the 10000 NCVs, corresponding to these components. Next, the fitted UMAP was used to embed all PCA-transformed NCVs to a 3D space. Finally, these 3D coordinates were re-normalized and encoded into a perceptually uniform CIELAB colorspace. Selecting molecules uniformly across multiple dimensions was done by taking sum over all PC coordinates for each of the molecules, ranking them by the obtained values, and then selecting a subset from this array uniformly across indices. Such approach allows to have a subset of molecules with density, similar to the original distribution while avoiding stochastic sampling. If assignment of molecules to background noise or true signal is available for a given dataset, only non-background molecules are used for fitting UMAP.

### Markov Random Field segmentation

In many cases, spatial proximity of molecules is a sign of their similarity by some other properties. Examples of such properties include cell assignment, cell type that produced the molecules, or the distinction between “background” and true “intracellular” molecules. To infer such labeling from molecules we formulate it as a segmentation problem, which is solved using an Expectation-Maximization (EM) algorithm for separating a mixture of distributions with Markov Random Field (MRF) prior^21,26^ to encode spatial relationships.

Such a model implies that each segment (which can be cell, cell type, or a background/signal label) comes from its own distribution that is generated by

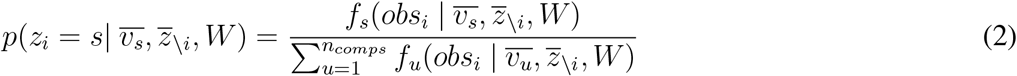

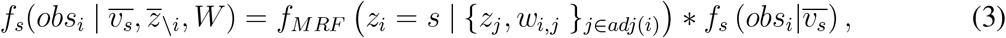

where *f_MRF_* is the MRF density, *f_s_* is the density of the component s, *n_comps_* is the total number of components, *z_i_* is the segment label for the molecule *i*, 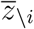 is the vector of labels for all molecules except i, *obs_i_* is the observed data from this molecule, *W* = {*w_i,j_*}_*i,j∈molecules*_ is the matrix of MRF edge weights between pairs of molecules *i* and *j*, and 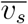 is the vector of parameters for the component *s*.

#### Building the Random Field

To establish the structure of the random field, Delaunay triangulation over points in 2D space was built using the VoronoiDelaunay.jl package. It provides a connected planar graph, matching the general structure of the space. Edge weights were set to the trimmed inverse Euclidean distance, so they represent connectivity of the two molecules *i* and 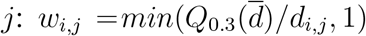, where 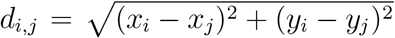, and 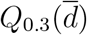 is the 0.3 quantile of the distance distribution 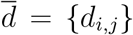. In principle, the weights could be additionally adjusted to represent other kind of dependencies between molecules (see the “Cell Segmentation” section), but they were not used during this step. It is worth noting that the Delaunay triangulation captures only a small neighborhood of a molecule, which is necessary to keep the graph planar. However, more neighbors could, in principle, be taken into account by adding edges to the nearest neighbors that are not already connected.

#### Separation of the intracellular molecules from the background

In the existing datasets, background regions are much sparser than the cellular somas. This difference can be quantified by estimating distance to the *k*-th nearest neighbour (NN) for each molecule. The molecules in the dense regions would have small distance, while for sparser regions the distance will be large (2c,d). To model distribution of such distances *d_i_* we used Gaussian Mixture Model with two components: one for the intracellular and another for background molecules. The EM algorithm with the MRF prior was then used to separate this components.

Initialization was performed using 10’th and 90’th percentile of the distance distribution for the means of the intracellular (*μ_c_*) and the background (*μ_b_*) components correspondingly. Both standard deviations were initialized as *σ_c_, σ_b_* = 0.25 ∗ (*μ_b_ − μ_c_*). The probability of a molecule to be labelled as intracellular (later called “molecule confidence” for simplicity) was initialized as 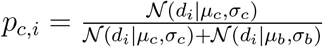. The molecules from the right tail with *d_i_ > μ_b_* + 3*σ* were assigned to the background with probabilities fixed to 1.0 and excluded from subsequent optimization. Finally, the total number of molecules per component was initialized as 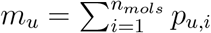.

On the Expectation step of the iteration *t* the assignment probabilities were updated using the following formulas with *u* ∈ [*c, b*]:

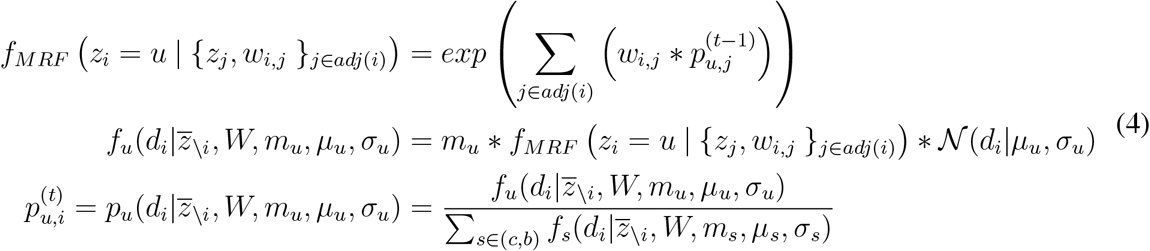

On the Maximization step parameters 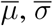 and 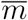 were re-estimated according to:

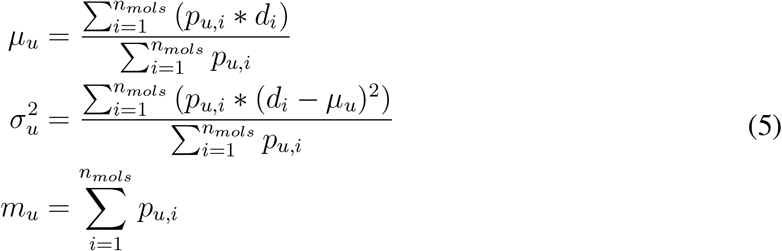

The difference of the parameters 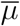 and 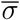 between iterations was used as the convergence criteria, with the convergence threshold of 0.005. After the algorithm converged, the MRF prior was discarded and only the densities of the normal distribution were used to estimate the cell assignment probabilities: 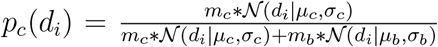. That was done because the MRF prior consistently push probabilities to be close to either 0.0 or 1.0, which corresponds to binary classification. In contrast, using only normal densities results in more gradual probability values, which can be integrated better with the further probabilistic parts of the algorithm.

When a prior segmentation *L_prior_* was available (*e.g*. DAPI), it was used as a constraint for the optimization. Given the prior segmentation confidence *c_prior_* ∈ [0.0,1.0], the Expectation step was restricted *p_c_*(*d_i_*) to be greater or equal than 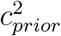 for all molecules, assigned to cells in the prior segmentation:

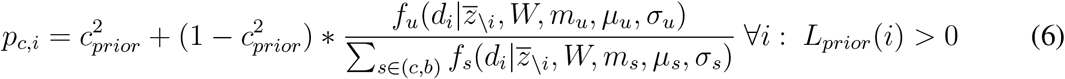

At the limit of *c_prior_* → 0, this approach converges to the case without a prior segmentation, whereas *c_prior_* → 1 will ensure that the molecules assigned to cells in the prior segmentation will be necessarily recognized as intracellular.

#### Segmentation of cell types

The same MRF formalism can be applied to perform de-novo clustering of molecules by cell type of origin. Moreover, when the scRNA-seq data is available, the approach can match the transcriptional identifies of the clusters identified in the spatial data to those observed in the provided scRNA-seq. To perform the segmentation, we considered gene identity of a transcript as observable data, and modeled the whole dataset as a mixture of Categorical distributions representing different cell types. The number of components *n_comps_* is an input algorithm parameter that must be specified at the start. Then, the basic Expectation step on the iteration t works as the following:

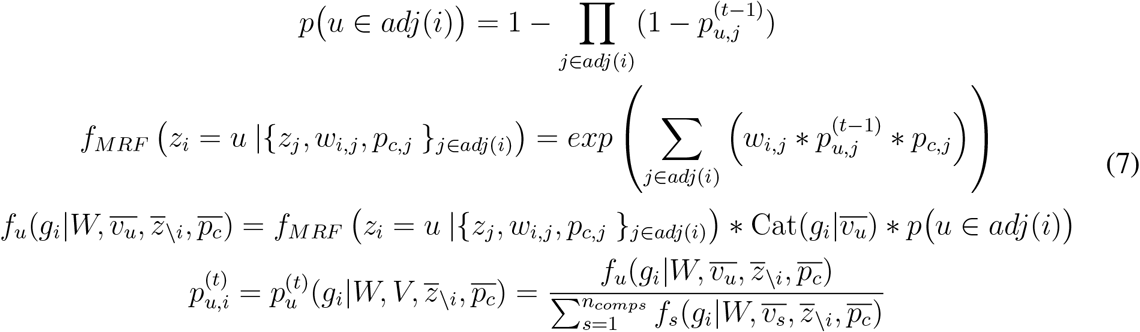

These formulas differ from the previous case (4) in several aspects. First, given that gene identities are categorical, and not continuous variables, the two Normal distributions were replaced with *n_comps_* Categorical ones to describe this kind of data. The parameter 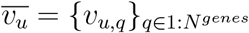 here defines expression probabilities per gene for component u. Second, they remove dependency of the density on the segment size *m_u_*, as its presence forced the algorithm to prefer components of larger size, and without strong signal from data, the MRF prior tended to eliminate all but the largest component. Such problems did not arise in the previous application, because there two Normal distri-butions described the observed data much better than one, ensuring that both components (background and signal) were maintained. In contrast, here the whole dataset can be described as a single Categorical distribution with a high quality of fit, pushing the algorithm to converge towards one component. Next, the molecule confidence *p_c,j_*, obtained from the background estimation step, is used here for estimation of *f_MRF_*. This step is very important, because only intracellular molecules are expected to be clustered over cell types, while background molecules can be arranged in arbitrary patterns. Finally, the updated formulas introduce *p*(*u ∈ adj*(*i*)), which ensures continuity of the resulting segmentation: without it, a molecule can be assigned to some cluster even if none of its neighbors belong to that cluster, as *f_MRF_* (*z_i_ = u* |{*z_j_, w_i,j_, p_c,j_* = 0}_*j∈adj*(*i*)_) = *exp*(0) = 1. Indeed, such ability to re-assign molecules globally can be desirable, as the EM algorithm can get trapped in local minima and is generally sensitive to initialization. And the assignments to the components distant from the local neighborhood can help to reach convergence to a global optimum. However, the probability of assigning a molecule to a cluster that is not connected to it cannot be inferred solely from the density *f_MRF_* (*z_i_ = u*) = 1; it also depends on the scale of the other terms. So, in addition to *p*(*u ∈ adj*(*i*)) the formulas introduce an additional term, *p_global_* = 0.05, which defines the probability of assigning the molecule to a component regardless of the connectivity. Then, the Expectation step can be adjusted as:

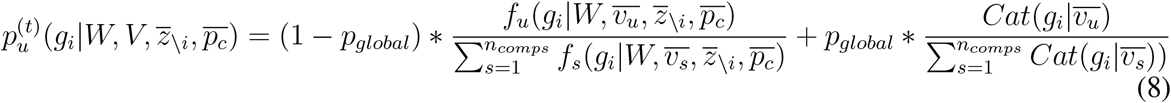

The downside of the global assignment is the reduced continuity of clusters. To correct for that effect, after the algorithm converged, *p_global_* was set to 0, which reduces to the basic version of the Expectation step, and the EM iterations were carried out further until convergence to the final result.

The Maximization step fit the parameters for 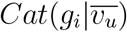 using the assignment *p_u,i_* from the Expectation step:

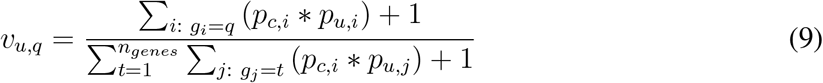

It is important to note here that the vector 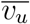 is a non-normalized probability density, due to the way pseudo-counts were incorporated: instead of adding *n_genes_* to the denominator, 1 was added as it allowed to preserve spiking structure of the sparse distribution.

When prior information about transcriptional composition of cell clusters is available (*e.g* scRNA-seq cell types), the Maximization step is changed to take it into account. Given prior expression fraction *μ_u,q_* from the cluster u and the gene *q* and its standard deviation *σ_u,q_*, the estimate was adjusted based on the Z-score value 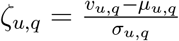 as:

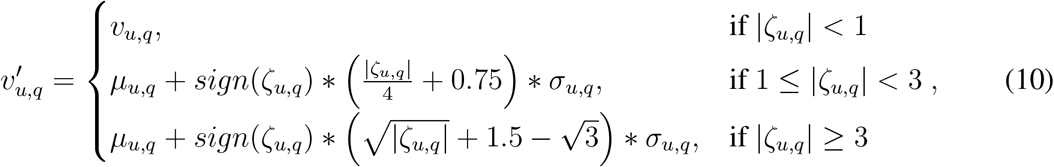

This function corresponds to no penalty for all deviation from the mean within one *σ*, linear penalty for deviations less than 3*σ*, and a super-linear penalty otherwise (Supplementary Fig. 14a). If the standard deviation was not available, it was set to the mean value by default: *σ_u,q_ = μ_u,q_*. It is important to note here that for a given prior clustering reasonable results can often be obtained by running only the Expectation step without the Maximization, which corresponds to setting *σ_u,q_* = 0, ∀_*u, q*_. Particularly, the results shown on the Supplementary Fig. 5 were obtained with only iterating over the Expectation step.

The algorithm was initialized using a vector of gene frequencies, estimated over the whole dataset, multiplied by uniform noise in [0.95; 1.05]: 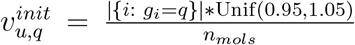. Next, these values were normalized by the sum over all genes: 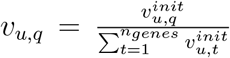. The initial assignment probabilities were estimated as 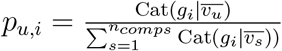. Convergence was determined based on the maximal change in *p_u,i_* between iterations, weighted by 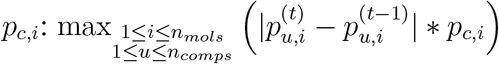.

The default threshold for the convergence was set to 0.01.

### Cell segmentation

Assignment of molecules to a cell of origin is another case of the MRF segmentation. In the most basic form, a cell can be modeled as a Multivariate Normal distribution over positions of molecules within a cell and a Categorical distribution over the cell gene composition. Thus, the density of a cell *k* in the molecule *i* is:

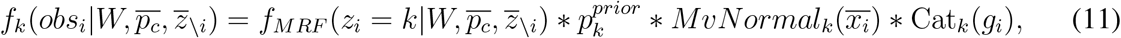

where 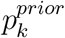 is the prior probability of the component *k*, which is proportional to the number of molecules, assigned to this component, 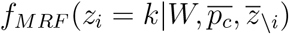 is the MRF term, and *MvNormal*() is the multivariate Normal distribution (bivariate for the 2D implementation). However, fitting this model by the EM algorithm does not work well for several reasons. First, the number of components cannot be defined beforehand and has to be inferred by the algorithm. To perform such inference, the algorithm we use a Dirichlet prior over the number of components (the approach called Bayesian Mixture Modeling or BMM). Second, not all of the molecules belong to cells, and filtration of the background molecules by a hard threshold over *p_c,i_* does not always work well. So the algorithm was adjusted to deal with raw probabilities *p_c,i_*, and a separate background component *f_bg_* was added to the mixture. Finally, dense homogeneous regions within a tissue do not have sufficient information to be segmented based on their transcritpional composition, so the model (11) will segment the whole region into a single component. Such situations, however, can be resolved much better by utilizing information about the expected physical size of a cell. This was done by introducing a global scale parameter *s_global_* that was used as a prior for estimating the covariance matrices of the bivariate Normal distribution. If a prior segmentation is provided, *s_global_* is inferred based on this data, and otherwise it has to be specified by the user.

#### Initialization

The algorithm is initialized with a large number of components uniformly distributed over 2D space. Setting the initial number of cells to be much larger than the expected number greatly improved the convergence of the algorithm. The initial cell centers were selected uniformly across 2D space, and each molecule was assigned to the nearest center. The center selection was done using a strategy similar the one used for subsampling of the neighbourhood composition vectors (NCVs): the molecules are ranked by the sum over the x and y coordinates, and then the algorithm selects a subset from this array uniformly across indices. When molecule background confidences were available, only molecules with true signal confidence greater than 0.25 are used. The MRF was initialized using Delaunay triangulation in the same way as described above. However, given that the triangulation uses only the information about spatial positions, the MRF was then further adjusted to reflect the neighbourhood composition similarities, based on NCVs. Specifically, NCVs were estimated for each molecule, and the resulting matrix of NCVs was transformed using PCA. Pairwise Pearson linear correlation of the PC vectors, *ρ_i,j_*, was estimated for any two molecules *i* and *j* that were connected by an edge in the Delaunay graph, and the MRF edge weight was then multiplied by *max*(*ρ_i,j_*, 0.01).

#### Fitting Bayesian Mixture Model

Fitting of the Bayesian Mixture Model was performed by incorporating Stick-breaking process into the Stochastic EM algorithm. The algorithm iterates over the four steps over a pre-defined number of iterations *N^iter^* (500 by default, which was enough for convergence on all of the tests):

1. Maximize parameters of the distributions given existing assignment of molecules by cells (Maximization step)
2. Sample empty components from the Dirichlet prior (Distribution Sampling step)
3. Stochastically assign molecules to components given the exiting components (Stochastic Expectation step)
4. Remove all components that have less than two molecules assigned to them After finishing the iterations, the algorithm re-estimates molecule assignment by averaging it over the last *N^est^* iterations (*N^iter^*/10 by default).

#### Maximization

The Maximization step fits the parameters of the Normal and Categorical distributions based on the current assignment of molecules to components. For the Categorical distribution of the component *k*, non-normalized probability *v_k,q_* of the gene *q* being expressed was estimated as a fraction of the gene *q* across the observed molecules (smoothed with pseudo-counts), weighted by the molecule confidence *p_c,i_*. To avoid numerical problems, all confidences were restricted with 0.01 from the bottom: 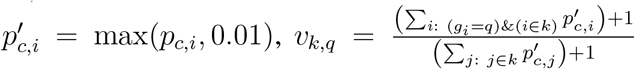. This procedure is similar to the one employed in estimation of probabilities for the cell type segmentation. However, here hard assignment of cell labels was used for the sake of performance. For the Multivariate Normal distribution, the mean 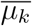 was estimated as the weighted average over the positions of the assigned molecules: 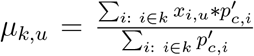, where *u* ∈ {1, 2}. The weighted Maximal Likelihood Estimator was also used for the covariance matrix: 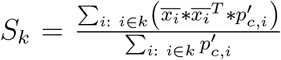. After estimating the covariance matrix, it was adjusted based on the global scale *s_global_*. The most popular solution for such adjustment uses the Wishart prior over the covariance matrix, as this prior is conjugate for the Normal distribution. However, the Wishart prior is parametrized with the expected covariance matrix and the number of degrees of freedom, and thus does not allow to explicitly control the magnitude of deviation from the expected covarate matrix. Therefore we instead relied on the non-conjugate Normal-like prior on the eigenvalues of the covariance matrix. This prior was parametrized with the expected size of the eigenvalues *s_global_*, and their standard deviation *σ_global_*, which can be specified by the user, or set to 0.25 ∗ *s_global_* by default. Then, the adjustment starts by performing eigen decomposition over *S_k_* to calculate its eigenvalues 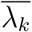 and the eigenvector matrix *Q_k_*. Next, the eigenvalues are adjusted using the formula 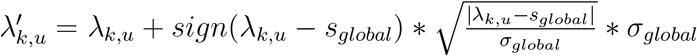, where *u* ∈ [1, 2]. This transformation corresponds to quadratic penalty over deviation Z-scores *Z_k,u_*, reducing the deviation *Z_k,u_ ∗ σ_global_* to 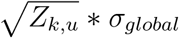. Next, to account for the components with low number of samples, it is further adjusted as 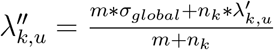, where *m* is the expected minimal number of molecules per cell and *n_k_* is the number of molecules assigned to the component *k*. Finally, the adjusted covariance matrix is estimated as 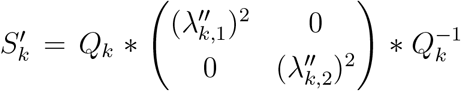. When the distribution parameters are estimated, the component prior probabilities 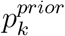 are sampled from the Dirichlet distribution, proportionally to the number of molecules per component 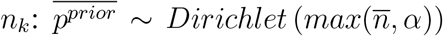, where *α* is the Dirichlet Process parameter, set to 0.2 by default. Density of the background component was also estimated during this step, as it is constant and depends only on the parameters of the cell components:

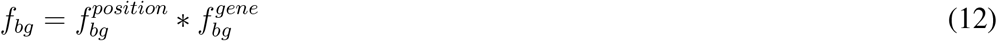

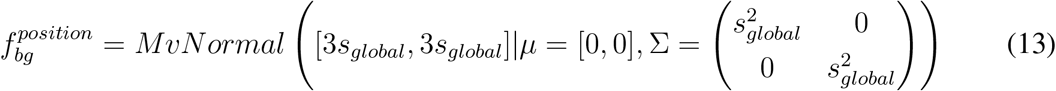

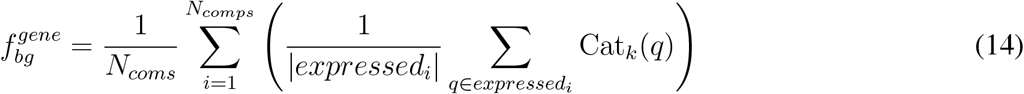

where *expressed_i_* is a set of all genes, expressed in the component *i*. The *position* part of this density corresponds to the level of three expected standard deviations (*i.e*. 3 ∗ *s_global_*) from the cell center. And the *composition* part estimates the average expression probability across all cells and all genes expressed in them.

#### Distribution sampling

After the Maximization step, *N_new comps_* = *β_new_ ∗ N_comps_* new components were sampled from the prior distributions. The parameter *β_new_* was set to 0.3 by default. For the Normal distribution, centers were sampled from all molecule positions with the weights, proportional to the molecule confidences: 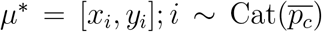. The diagonal covariance matrix was sampled from the global scale prior: 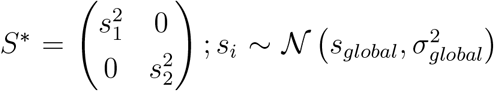. Sampling gene composition parameters requires proper modeling of sparsity of the expression vectors, which varies greatly between protocols and cell types. It is therefore unclear how to capture this with a parametric prior distribution. Instead, the algorithm sampled gene composition parameters uniformly from the existing components: 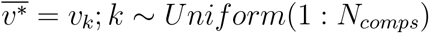.

#### Stochastic Expectation

During the Expectation step, the algorithm iterates over all the molecules and stochastically re-estimates their component assignment. The algorithm starts by determining candidate components for each molecule, as well as their MRF prior densities. First, if a component *k* already has some molecules {*i*}_*z_i_=k*_ assigned to it, it is included as a candidate for all molecules *j* that are connected to the assigned molecules {*i*}_*z_i_=k*_: (*z_i_ = k*) ⟹ (*k ∈ candidates_j_*, ∀_*j*_ ∈ *adj* (*i*)). The MRF prior densities are defined as the weighted sum over all edges coming from the molecules that are assigned to this component: 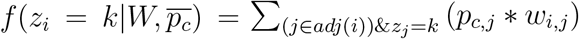. For every component *k* that was just sampled from the prior and still has no assigned molecules, the algorithms found the molecule *i*, closest to the component center 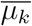 and included component *k* as a candidate for all molecules *j* that are connected to *i*:

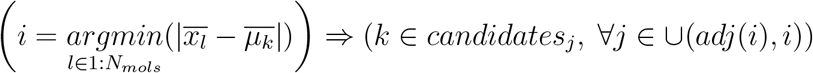

In this case, the edge weight was set to 1, which is the maximal possible weight given the scaling procedure. The background component was included as a candidate to all molecules, and its MRF prior probability was set to

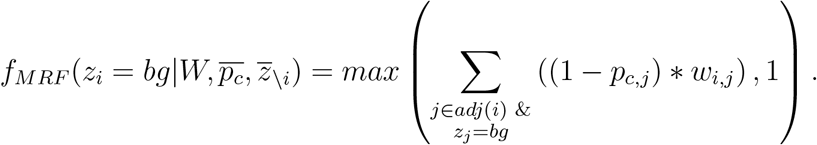

It’s important to note that the exponent was not used for the MRF densities here, as it gave visibly worse results on the tests. After the MRF weights are estimated, the full formula for the component density is:

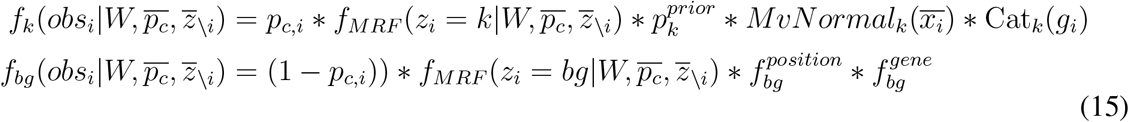

Finally, assignment for the molecule *i* is sampled from the estimated densities.

#### Using molecule cell type segmentation information

To improve gene composition purity of the cell segmentation, results from the cell type segmentation can be utilised here by penalizing assignment of molecules from different clusters to the same cell. For that, the cell type segmentation is ran prior to the cell segmentation, and each molecule has a cluster label assigned to it. By default, number of clusters is set to 4, as almost every dataset has 4 cell types, while having larger number of clusters could lead to over-segmentation. Given a cluster label per molecule, on the maximization step the algorithm estimates the most represented cluster per cell. Then, on the expectation step, densities of the cell components are penalized in case the cluster of the molecule *cl_i_* does not match to the cluster *cl_k_* of the cell: 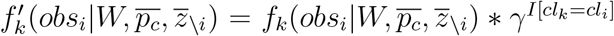, where *γ* is the penalty term (0.25 by default) and *I* is the indicator function.

#### Using a prior segmentation

For many datasets, the auxiliary microscopy stains, such as DAPI, can be used to generate a prior segmentation. Such a segmentation can be used to resolve the cases where molecules do not provide sufficient information. The complexities of segmenting and aligning the auxiliary stains, however, mean that the quality of such segmentations can vary. To incorporate this optional information, Baysor can accept prior cell segmentation labels per molecule, as well as the prior segmentation confidence parameter *c_prior_* ∈ [0; 1]. At *c_prior_* = 0, the prior segmentation would be ignored, whereas *c_prior_* = 1 forces the Baysor segmentation not to violate the prior segmentation assignments. Increasing *c_prior_* from 0 to 1 gradually increases importance of the prior segmentation. The prior segmentation penalty is evaluated only for the molecules that are assigned to some cell (but not to background) in both Baysor and the prior segmentations. This accounts for the fact that the imaging-based segmentations may miss some cells or portions of cells that can still be deduced from the spatial transcriptomics data. The most obvious example of such situation are the DAPI-based segmentations, which cover only molecules within the cell nuclei, leaving most of the cytoplasm molecules unannotated. Baysor algorithm, therefore, treats background labels (*i.e*. not within segmented region) within the prior segmentations as “unknown”, instead of explicitly assigning these molecules to the background component. The situation in which some non-background molecules from the prior segmentation are recognized as background by Baysor is dealt with during the Background segmentation step. Specifically, if a molecule *i* is recognized as intracellular in the prior, its confidence can not be less than 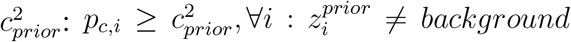. As a result, there are two possible types of contradictions between the two segmentations: (i) multiple Baysor components are present within one prior segment, and (ii) one Baysor cell component touches multiple prior segment.

The penalties are applied during the stage at which the densities of components are estimated for a given molecule, by multiplying 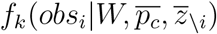 by the penalty term *β_k,i_*. If this molecule has a prior assignment to some segment, all but the component with the largest intersection with this prior segment are penalized. This is done using the following procedure. After the Distribution Sampling step, the algorithm estimates number of molecules 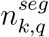 per segment *q* for each of the cell components *k*. Then, based on these numbers it estimates the main prior segment per component *ς_k_*. This main segment label is used to separate the cases where two Baysor components are present within one prior segment (in this case they would have the same main segment id) from the cases where a Baysor component touches multiple segments (it would have one main segment, but would get penalized for touching all other segments). During the Expectation step for the molecule *i*, the algorithm finds the component *u*\*, which has the largest intersection with the segment 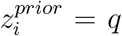 across all candidate components for any molecule *i* that has this segment as their main assignment or do not have a main segment at all: 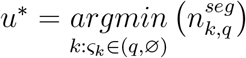. If any two components have the same number of molecules for this segment, the component with larger total number of molecules is chosen as the main component. The main component per segment incurs no penalty (*β_u*,i_* = 1), while the others are penalized as the following:

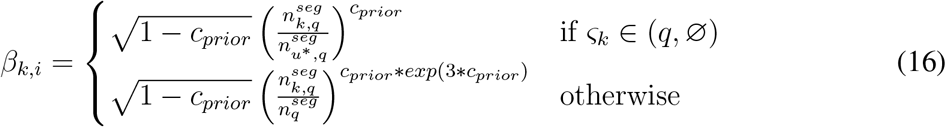

where 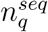 is the total number of molecules per segment *q*. The first type of penalty prevents two cells from existing in the same prior segment (over-segmentation problem), while the second type prevents one cell from taking over a big part of a segment assigned to a different cell (overlapping problem). The second penalty type also solves the under-segmentation problem: if a Baysor cell overlaps two prior segments, then as soon as a new component is sampled inside one of these segments, the existing “doublet” cell incurs very large penalty for all other molecules of this segment. These functional form of the penalties, while somewhat arbitrary, has allowed to express a desired curve shape for the penalties (see Supplementary Fig. 14b,c). The penalties are also scaled in [0; 1], which allows to keep the Distribution Sampling step without changes. Freshly sampled cells have no penalty, which corresponds to *β_k,i_* = 1.

#### Complete Baysor workflow

Below is the description of the full Baysor workflow, as implemented by the command-line interface:

1. Read the data frame with information about molecules. Filter out the genes with total number of molecules below threshold if it was specified.
2. If provided, load the prior segmentation mask. Filter out any prior segments that have less than *m^prior^* molecules. Estimate the global scale *s^global^* based on this data.
3. Estimate confidence *p_c,i_* per molecule
4. If the specified number of cell clusters (4 by default) is greater than 1, run the cell type segmentation using the confidences, estimated above. Otherwise, assign all molecules to the same cluster.
5. Initialize the Bayesian Mixture Model algorithm
6. Optionally, split data by frames to enable parallel processing
7. Run the BMM algorithm
8. Re-estimate assignment by averaging over the last *N^est^* iterations
9. If the data was split by frames, merge them back into one
10. Save the segmentation results

#### Inferring the algorithm parameters

Baysor implementation derives most of the parameter estimates based on the minimal expected number of molecules per cell *m*, which must be specified by the user, as well as the number of genes measured by the assay (*n_genes_*):

- The initial number of cells is set to 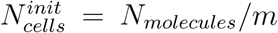, where *N_molecules_* is the total number of molecules.
- The number of principal components for the PCA transformation of NCVs for adjusting the MRF is estimated as min (max (*n_genes_*/3, 30), 100, *n_genes_*)
- The nearest neighbor index for estimating distances *d_i_* during the Background Segmentation is set to max (*m*/2 + 1,2)
- The number of nearest neighbors for NCV coloring is set to max(*n_genes_*/10), *m*, 3)
- If a segmentation mask was provided, it is used to infer the global scale *s_global_* and its standard deviation *σ_global_*. To do so, Baysor first approximates cell radii from the number of pixels per prior segment 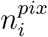 as 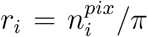. Then, 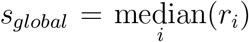 and 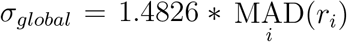, which are the median and the adjusted median absolute deviation over the radii.
- Without a segmentation mask, the global scale *s_global_* is a required input parameter, and the deviation *σ_global_* is set to 0.25 ∗ *s_global_* by default.
- The minimal number of molecules per prior segment *m^prior^* = max(*m*/4, 2).

#### Segmentation parameters

Parameters of the segmentation runs for different datasets are shown in the Supplementary Table 3.

### Benchmarks

To evaluate Baysor, we have shown its performance in separating distinct cell types, using Pearson Correlation benchmarks. We have also shown stability of its convergence for different random number generator seeds, as well as runtime profiling.

#### Correlation benchmarks

As we currently lack independent techniques to establish ground truth on cell segmentations, we developed a benchmark to evaluate the differences between two segmentations. For each source cell *c* in the segmentation 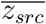 the benchmark finds a target cell *t*\* from the segmentation 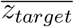 that has the largest overlap with *c* (Fig. 4c). Then it splits the molecules from *c* over the part that overlaps with *t*\* from the molecules in the non-overlapping part. Measuring similarities of gene compositions of these two parts as a Pearson linear correlation (*ρ_c_*), the benchmark evaluates the distribution of such similarities over all cells from the source segmentation: 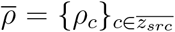. Taking two segmentations 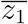 and 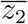 as the input, the benchmark returns two distributions 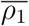 and 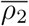 for the cases where 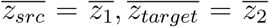 and 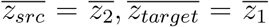 correspondingly. Based on these two distribution, the segmentation is said to be better if it has higher 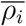 values. More formally, given two segmentations 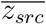 and 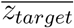, the benchmark procedure does the following:

1. All cells *i* with the total number of molecules below the threshold 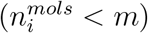 are removed from both segmentations.
2. Among the remaining cells, a contingency matrix between the two segmentation assignments is estimated.
3. Based on the contingency matrix, for each cell *c* of the segmentation 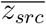 and cell *t* from the segmentation 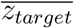, the overlap fraction 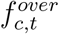 is estimated.
4. For each cell 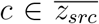, the target cell 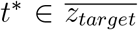 is determined as the cell with the highest molecule overlap fraction 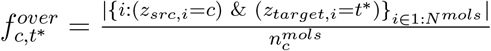.
5. Only cells from 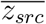 with the overlap 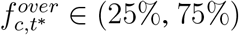 are selected for further analysis.
6. For each cell 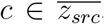, the molecules from this cell are partitioned into (i) the main overlap 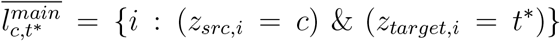, and (ii) the rest 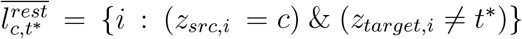.
7. Expression vectors 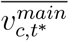 and 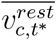 are estimated over the molecules from 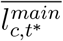 and 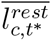 correspondingly.
8. Pearson linear correlation coefficients *ρ_c_* between the vectors 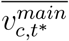 and 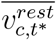 are estimated. If these two sets of molecules were produced by different cell types, the correlation *ρ_c_* is expected to be low, suggesting that the segmentation 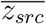 was erroneous for the cell *c*. However, similarly to the classical hypothesis testing framework, a high correlation *ρ_c_* does not mean that the segmentation 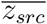 is correct: it only suggests that the molecules were obtained from the same cell type, and it is still possible that they originated from different cells.

#### Stability benchmarks

To assess the stability of the algorithm convergence, Baysor was run 10 times with different random number generator seeds on the MERFISH dataset. Each run used 400 iterations each, with parameters “*scale=6.16*” and “*min_molecules_per_cell=30*”. The final assignment was obtained by averaging over the last 100 iterations (“*assignment_history_depth=100*”). The molecule confidences and clusters were pre-estimated only once, as these procedures are deterministic. Pairwise Adjusted Rand Index and Mutual Information between results of different runs were calculated as stability metrics.

#### Performance benchmarks

For runtime performance profiling of the molecule clustering (Supplementary Table 1), MERFISH dataset was used with the molecules subset to those with coordinates *x* < −3300 and *y* < −3300 (338023 molecules in total). Then, molecule confidence was estimated, using *confidence_nn_id=26* parameter setting. The MRF clustering was then ran five times for each value of *k* ∈ {2,4,6,8,10}, with parameters *max_iters=5000* and *n_iters_without_up-date=100*.

To evaluate the performance of the NCV Leiden clustering, Pagoda2 package was used. For each run, the NCV matrix was estimated using 50 nearest neighbors, normalized by the total count of each NCV, and reduced to projections on the top 50 principal components. A k-NN graph was then built using 30 nearest neighbors (using the default cosine similarity distance metric) and leiden community detection algorithm was applied with the default *resolution=1.0*. For the profiling, the same parameters as specified in the Supplementary Table 3 (Prior=No) were used. Each dataset was segmented 5 times using a single thread. All benchmarks were ran on Lenovo Thinkpad X1 laptop with Intel(R) Core(TM) i7-8850H @ 2.60GHz CPU and 64GB RAM.

### Quality metrics

After finishing the segmentation, Baysor extracts the following summary metrics per cell:

- Cell area: area of the convex hull around the cell molecules
- Density: cell area divided by the number of molecules in cell
- Elongation: ratio of the two eigenvalues of the cell covariance matrix
- Average confidence: average confidence of the cell molecules

### scRNA-seq processing

#### Analyzing NCVs using scRNA-seq tools

To generate clustering and embedding of NCVs (Figure 1b,c) Pagoda2 package (https://github.com/kharchenkolab/pagoda2/) was used. The data was pre-processed using the total count normalization, and then building 50-NN graph over the normalized counts using the default cosine distance metric. Leiden clustering with parameters “resolution=8” and “n.iterations=15” was used for annotation expansion (see below). Visualization of the MERFISH and the osm-FISH datasets was done by building a joint matrix over the paper and the Baysor segmentations, reducing its dimensionality with PCA to 10 top PCs (MultivariateStats.jl package) and then running UMAP embedding (UMAP.jl package) using Euclidean distance and parameters “*spread=2.0*” and “*min_dist=0.1*” over the joint matrix. Hierarchical clustering with the Ward linkage method and 70 clusters (Clustering.jl package) was used for the annotation expansion.

#### scRNA-seq Annotation

The annotation for Allen smFISH data was generated using CellAnnotatoR package. The cell type markers were inferred from Mouse VISp scRNA-seq data produced for the SpaceTx consortium by the Allen Brain Institute (personal communication), and then applied it to the corresponding FISH data. For inference, the “merged_cluster_smFISH” annotation shared with the dataset was used. First, the cell type hierarchy was built using “broad_class” as the first level, class prefix from “merged_cluster_smFISH” as the second and the full “merged_cluster_smFISH” labels as the third level of the hierarchy. Second, cell types “CR”, “Astro”, “Endo”, “Macrophage”, “Oligo” and “SMC” were removed, as they could not be distinguished by the genes, measured in the Allen smFISH dataset. Next, CellAnnotatoR marker inference procedure was utilized to obtain the markers using this hierarchy, and the resulting marker list was adjusted by hands to improve the quality of the annotation. Finally, CellAnnotatoR was used to apply the identified markers to the NCV data. To annotate MERSISH and osm-FISH data, the marker list was compiled manually using the information published by the protocol authors, and then CellAnnotatoR was applied in the same manner. In all three cases, the resulting annotation was expanded over the clusters found in the previous step (see “Analyzing NCVs using scRNA-seq tools”).

#### Polygon visualization

To visualize the boundaries of the cells, a Kernel Density Estimation (KDE) based algorithm was used. The algorithm builds a grid over 2D space, assigns each grid node to the cell with the highest density of molecules around this node and draws a polygon around this labels on the grid. More formally:

1. A uniform 4-connected grid was created over the 2D space, with the grid step specified as an input parameter.
2. The density of molecules for each cell was estimated over the grid nodes using the KDE implementation in the KernelDensity.jl package. KDE bandwidth - a parameter of the algorithm - was set to the 0.5 * “grid step” by default. To improve runtime performance, only the nodes within three bandwidths of the cell molecules were taken into account.
3. Each node was assigned to the cell with the maximal density in this node. If the maximal density was below threshold (10^−5^ by default), the node was assigned to the background.
4. For each cell, the graph of boundary nodes were determined as the grid nodes of the cell that were adjacent to the nodes from the other cells or from the background. The edges between these boundary nodes were accepted into the boundary graph.
5. For each cell, a minimal spanning tree was built over its boundary graph using the Kruskal algorithm.
6. For each cell, the longest path in this tree was extracted using the Dijkstra algorithm, and the resulting path was transformed into a polygon by connecting its beginning and its end.

#### DAPI watershed segmentation with ImageJ

To generate prior segmentations using DAPI stains, the Watershed segmentation was performed using ImageJ^23^ software. Each staining was segmented by the following procedure:

1. The image was converted to 8-bit format (“Image” / “Type” / “8-bit”).
2. Median filter with one pixel radius was applied (“Process” / “Filters” / “Median…”).
3. Auto Threshold with the “Default” method and “Ignore black” option was applied for image binarization (“Image” / “Adjust” / “Auto Threshold”)
4. Watershed segmentation was applied (“Process” / “Binary” / “Watershed”).

#### VISP multiplexed smFISH data generation

Multiplexed smFISH data of mouse primary visual cortex (VISp) was generated as part of the SpaceTx consortium. Tissue processing was carried out as previously described^27^, with some modifications. The description is taken from^19^.

Silanizationof coverslips (#1.5, Thorlabs CG15KH) was performed by plasma cleaning for 30 min in a Plasma-Prep III (SPI11050-AB), followed by vapor deposition of 3-aminopropyltriethoxysilane (APES, Sigma A3648) in a vacuum for 10 minutes. Coverslips were then washedin 100% methanol for 2 × 5 minutes, allowed to dry, and stored in a dust-free environment until use.

Fresh-frozen mouse brain tissue was sectioned at 10 *μ*m onto silanized coverslips, let dry for 20 min at −20°C, then fixed for 15 min at 4°C in 4% PFA inPBS. Sections were washed 3 × 10 min in PBS, then permeabilized and dehydrated with chilled 100% methanol at −20°C for 10 min and allowed to dry. Sections were stored at −80 °C until use. Frozen sections were rehydrated in 2X SSC (Sigma 20XSSC, 15557036) for 5 min, then treated 10 min with 8% SDS (Sigma 724255) in PBS at room temperature. Sections were washed 5 times in 2X SSC. Sections were then incubated in hybridization buffer (10% Formamide (v/v, Sigma 4650), 10% dextran sulfate (w/v, Sigma D8906), 200*μ*g/mL BSA (ThermoFisher AM2616), 2 mM ribonucleoside vanadyl complex (New England Biolabs S1402S), 1 mg/ml tRNA (Sigma 10109541001) in 2X SSC) for 5 min at 37°C. Probes were diluted in hybridization buffer at a concentration of 250 nM and hybridized at 37^°^C for 2 h. Following hybridization, sections were washed 2 × 10 min at 37°C in wash buffer (2X SSC, 20% Formamide), and 1 × 10 min in wash buffer with 5 *μ*g/ml DAPI (Sigma 32670), then washed 3 times with 2X SSC. Sections were then imaged in Imaging buffer(20 mM Tris-HCl pH 8, 50 mM NaCl, 0.8% glucose (Sigma G8270), 30 U/ml pyranose oxidase (Sigma P4234), 50 *μ*g/ml catalase (Abcam ab219092). Following imaging, sections were incubated 3 × 10 min in stripping buffer (65% formamide, 2X SSC) at 30°C to remove hybridization probes from the first round. Sections were then washed in 2X SSC for 3 × 5 min at room temperature before repeating the hybridization procedure.

The multiplexed smFISH image data was collected and processed using methods previously described^27^, except that the images from different rounds of hybridization were registered in (x,y) based on the DAPI signal. The spot locations and raw data are available on request.

**Code availability.** Baysor package is available at https://github.com/kharchenkolab/Baysor. Baysor parameters for different datasets are reported in the Supplementary Table 3. The code to reproduce the results is available at https://github.com/kharchenkolab/BaysorAnalysis/.

