## Supplementary Figures and Table Legends for "Bayesian segmentation of spatially resolved transcriptomics data"

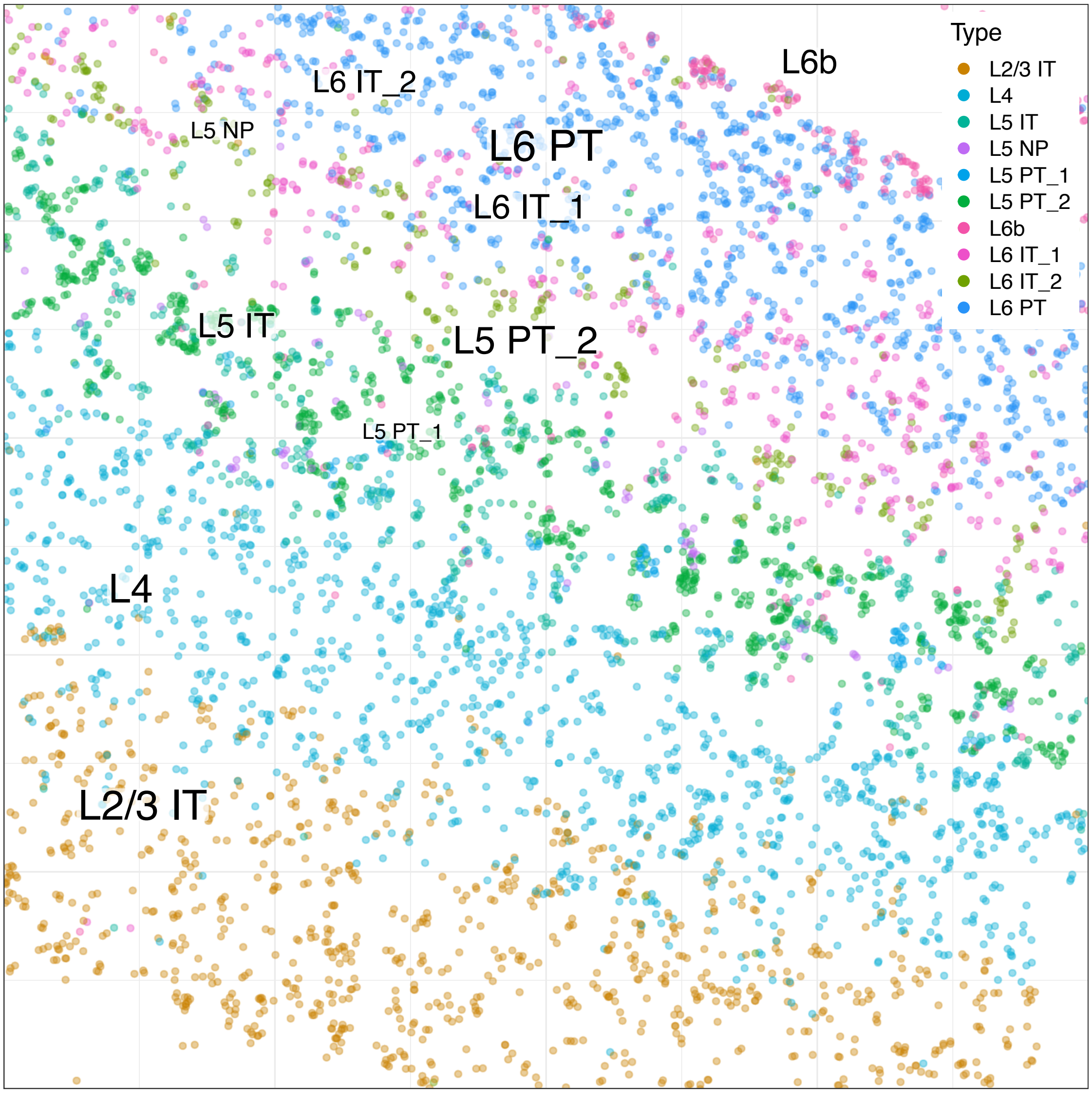


**Supplementary Fig. 1 | Visualisation of the layer structure on Allen smFISH dataset with Composition Neighbourhood Vectors.** The figure shows a subset of the Allen smFISH dataset. Each point represents a molecule in the physical 2D coordinates, colored by the annotated cell type of its Neighbourhood Composition Vector (see Fig. 1a) with neighbourhood size $k = 50$. Only a uniform subsampling of the molecules annotated as Excitatory neurons is shown.

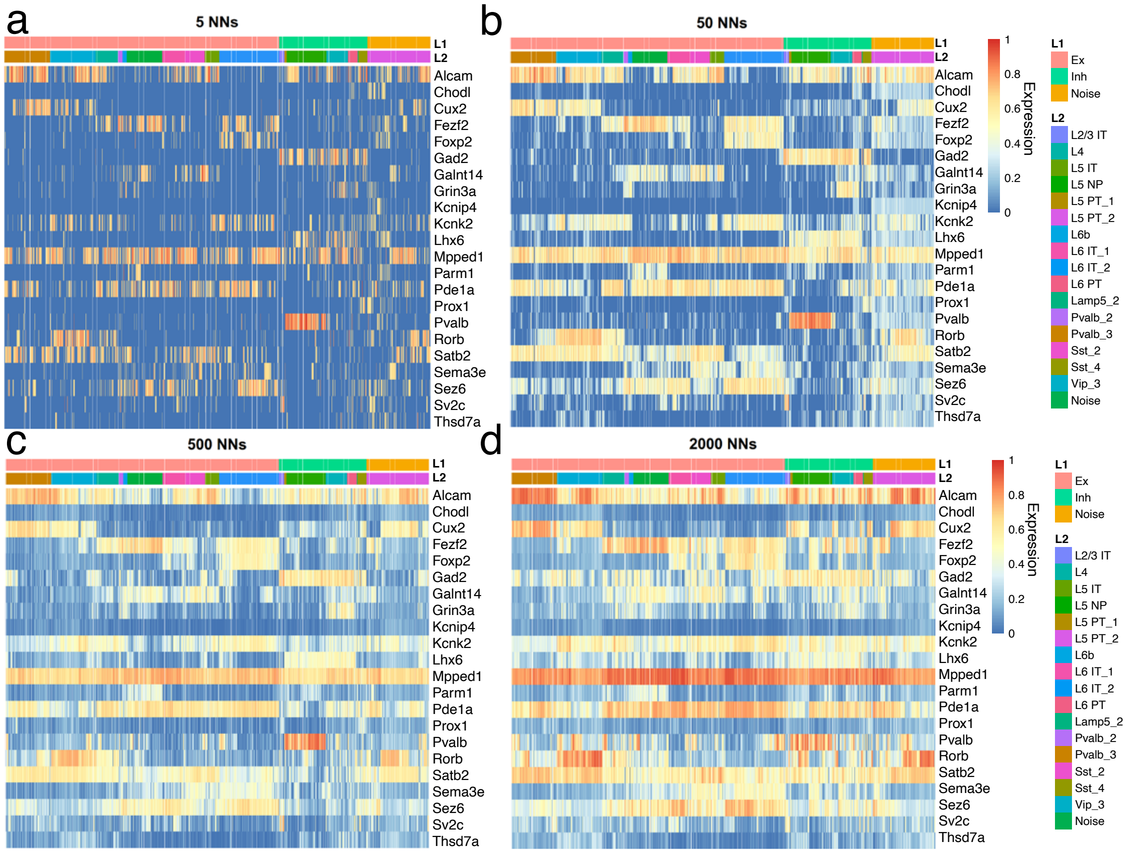


**Supplementary Fig. 2 | Expression patterns of the Neighbourhood Composition Vectors as a function of the neighbourhood size.** The panel shows expression patterns of the NCVs for the 20k molecules uniformly selected across the 2D space in the Allen smFISH dataset. These molecules were annotated based on 50-NN NCVs (Fig. 1b). The NCVs were then re-estimated using different neighbourhood size (shown in the plot titles). For each NCV, the expression vector was normalized by the total count and log-transformed. The whole expression matrix was then rescaled into the [0; 1] interval, using min-max scaling over the whole matrix. The heatmap color shows the scaled expression magnitudes, with the columns corresponding to the NCVs, and the rows corresponding to genes. The columns are ordered by the annotated cell type, and subsequently by the hierarchical clustering order within each cell type. It can be seen that low number of nearest neighbors (*i.e.* **a**) leads to a sparse NCV matrix in which the general patterns cannot be seen. On the opposite extreme, using very large number of neighbors (**c-d**) obscures expression patterns of individual subtypes.

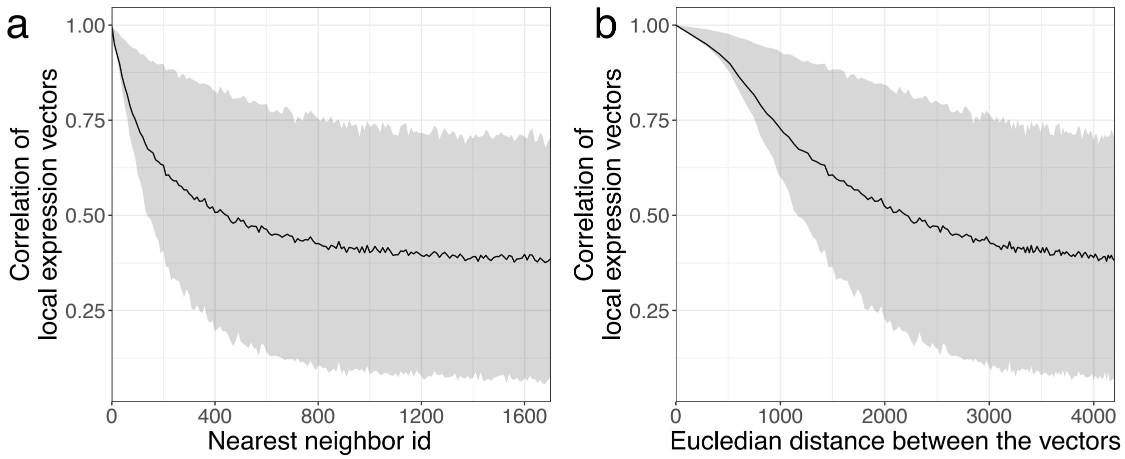


**Supplementary Fig. 3 | Pearson correlation of Neighbourhood Composition Vectors as a function of the distance between vectors.** The plots show the correlation of NCVs as a function of the nearest neighbor rank (**a**), or physical distance (**b**) for the osm-FISH dataset. The same 20k molecules were used as in the Supplementary Fig. 2. Neighbourhood size $k = 40$ was used. The mean values of the correlation coefficient are shown by a black line, and the gray area shows the 25'th and 75'th percentiles.

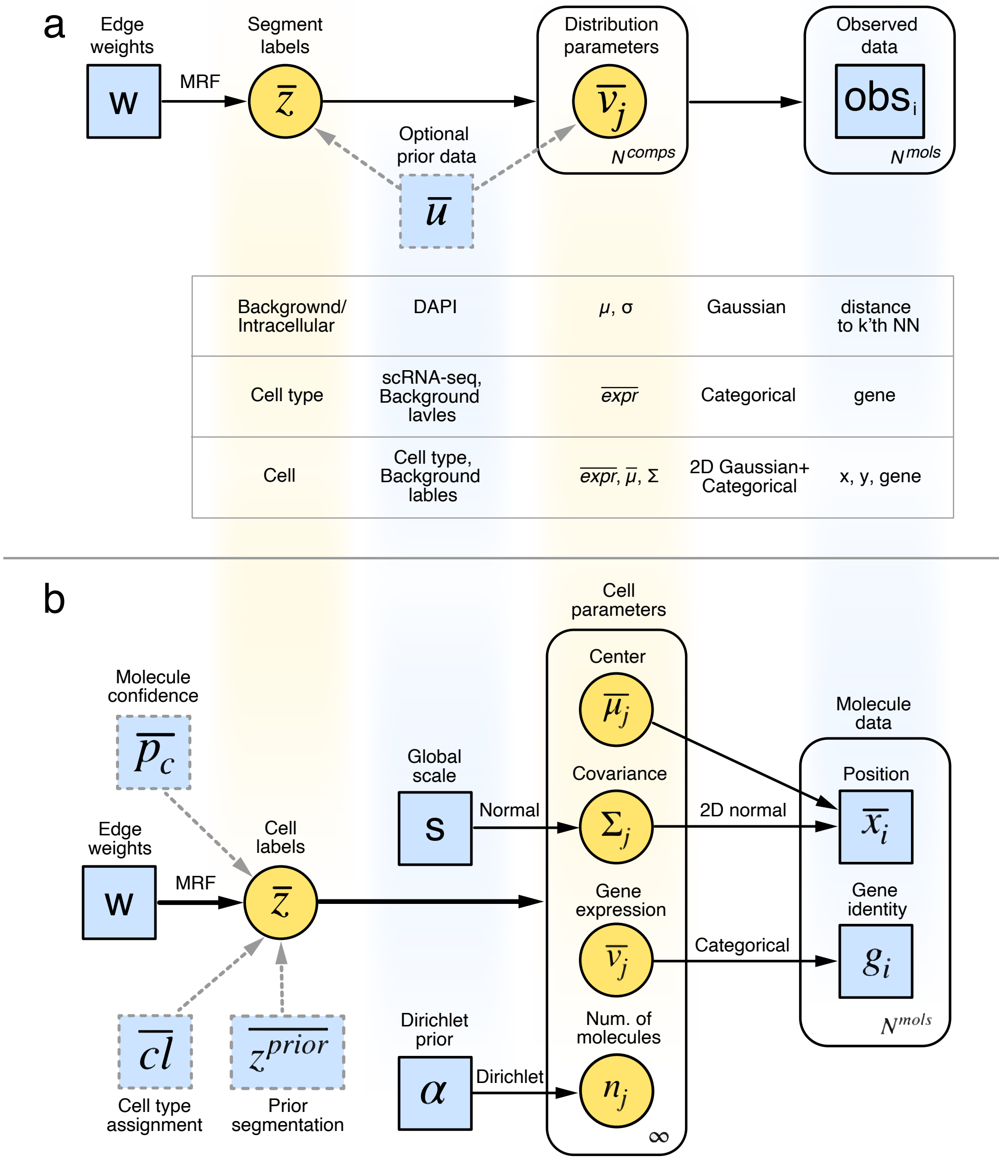


**Supplementary Fig. 4 | Graphical models of the segmentation process.** Graphical representations of the Bayesian models used for the general Markov-Random Field (MRF) segmentation process (**a**) and the extended model for cell segmentation (**b**) are shown. Blue squares represent input parameters and data for the algorithm. Optional input is shown with dashed border lines. The yellow circles represent the hidden parameters, fitted by the algorithm. Round-corner boxes represent plate notation for a mixture of distributions with the size of the mixture shown on the bottom right corner. $N^{mols}$ denotes the number of molecules in the dataset, and $N^{comps}$ is the specified number of the mixture components. Arrow labels show the distributions used to model dependencies between the corresponding variables. Matrix variables are shown with the capital letters and vector variables are designated with the overline. **a**, The general MRF model, where the MRF prior with weights **W** is used to account for the spatial dependency of the inferred labels $\overline{z} \in 1 : N^{comps}$. Examples of the variables and distributions for different labelling problems are noted below the boxes. **b**, The detailed model for the Cell Segmentation problem. Here, Bayesian Mixture Models with Dirichlet prior were used, so the possible number of components of the mixture is infinite, which allows the algorithm to estimate the number of components automatically. To ensure that the components correspond to the actual cells, the Global Scale parameter $s$ was introduced, which specifies the expected cell radius.

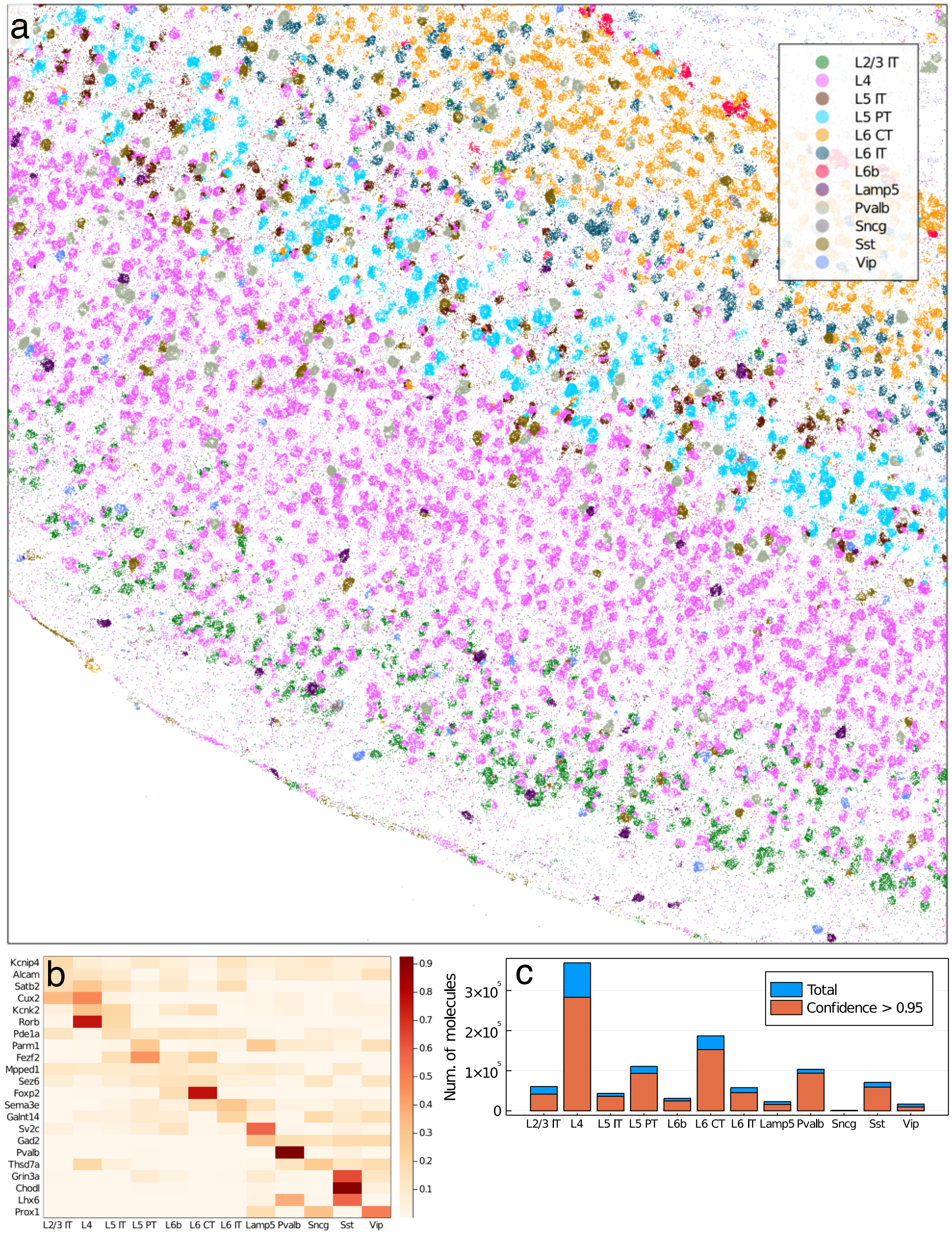


**Supplementary Fig. 5 | Annotation transfer using Markov Random Field segmentation with scRNA-seq prior.** Cell type centroids from the unpublished Mouse VISp dataset were estimated and used as a prior for MRF-based clustering of molecules from the Allen sm-FISH dataset. **a**, Cell type assignment is shown with color for each individual molecule, on a subset of the data in physical 2D coordinates. **b**, Average expression patterns of different genes (rows) are shown for the cell types (columns), as determined from the spatial data. The colors show the expression magnitude, normalised for each gene. **c**, The number of molecules (y-axis) per cell type (x-axis). The blue bars show the total number of molecules, with the orange bars representing only molecules with high probability ($> 0.95$) of being intracellular (as reported by the background segmentation algorithm).

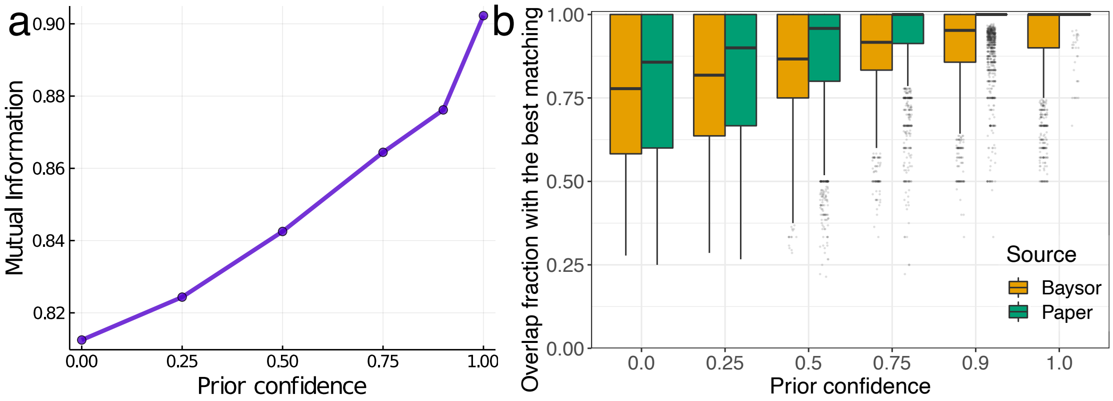


**Supplementary Fig. 6 | Impact of the "prior segmentation confidence" parameter on the difference between the prior and the posterior segmentations on the example of ISS CA1 region.** **a**, The Mutual Information between the Baysor and the Paper segmentations (y-axis) is shown as a function of the prior segmentation confidence (x-axis). Mutual Information does not reach the value of 1.0, as even for prior confidence set to 1.0, Baysor is still allowed to re-assign molecules, recognised as background in the Paper segmentation. **b**, For each cell of the source segmentation (shown with colour), a cell with the largest overlap was picked from the target segmentation. The overlap fraction is shown on the y-axis for the different values of prior segmentation confidence. It can be seen that for high values of the prior confidence, for each Paper cell there is a Baysor cell that covers it completely (confidence $\geq 0.9$, Source=Paper). The opposite is not true, as Baysor is allowed to re-assign the background molecules from the Paper segmentation.

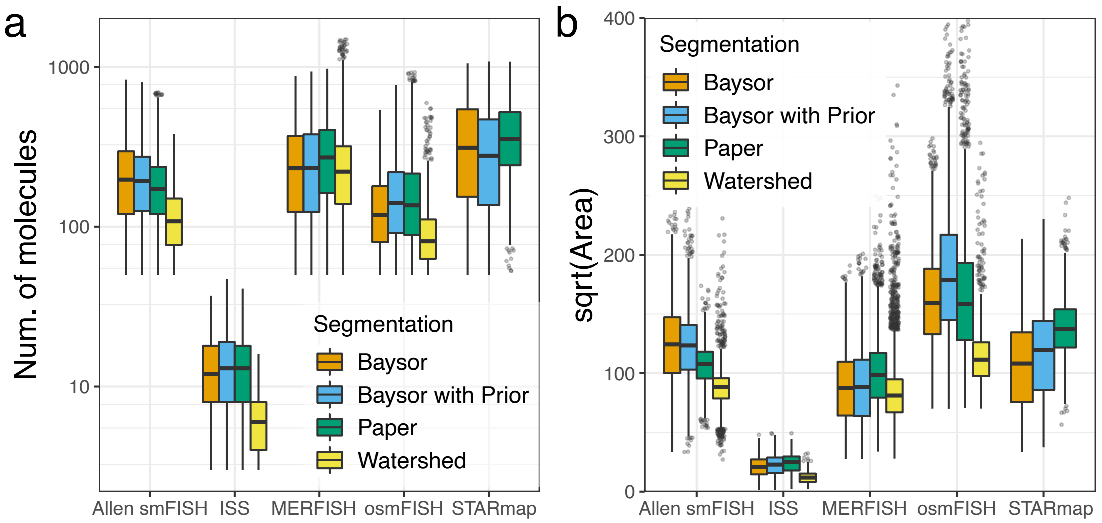


**Supplementary Fig. 7 | Cell statistics for different segmentation methods.** The boxplots show distributions of the number of molecules per cell (**a**, log-scale y-axis) and the squared root of the cell area, which is an approximation for cell radii (**b**) for different protocols (x-axis) and segmentation methods (fill colours). For all datasets, Baysor has approximately the same values as the published segmentations, which suggests that it is not biased towards over- or under-segmentation. The Watershed method stably shows lower values, which is expected as it only registering nuclei information while ignoring the cytoplasm molecules.

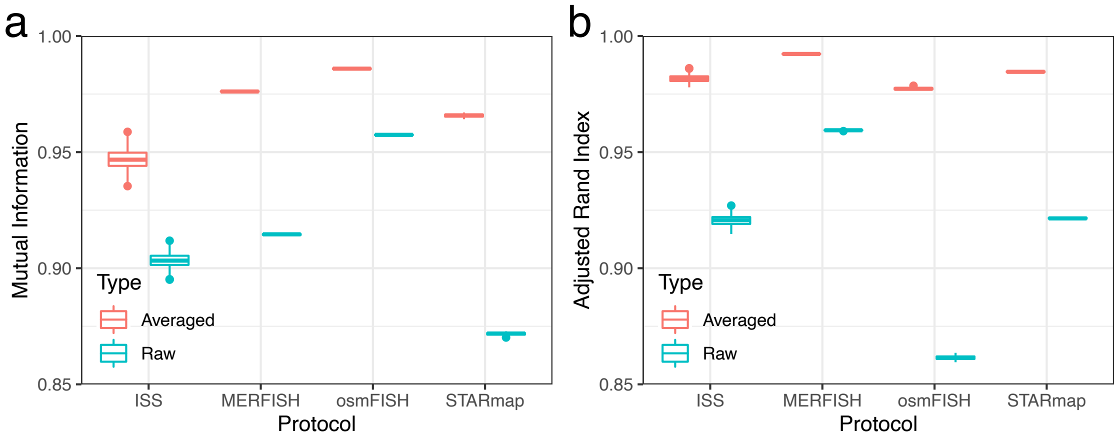


**Supplementary Fig. 8 | Cell segmentation stability.** The panels show stability benchmarks for Baysor cell segmentation, ran with the same parameters on the same data, but using different random number generator seeds. The segmentation was ran 10 times with different seeds, and the Mutual Information (**a**) and the Adjusted Rand Index (**b**) were estimated for segmentation runs. The benchmark was repeated for each protocol (x-axis). For better convergence, Baysor averages cell assignment across the last $N$ iterations of the algorithm ($N = 100$ shown in red). This strategy (red boxplots, left) strongly improves the stability compared to the naive approach of taking the assignment only from the last iteration (green).

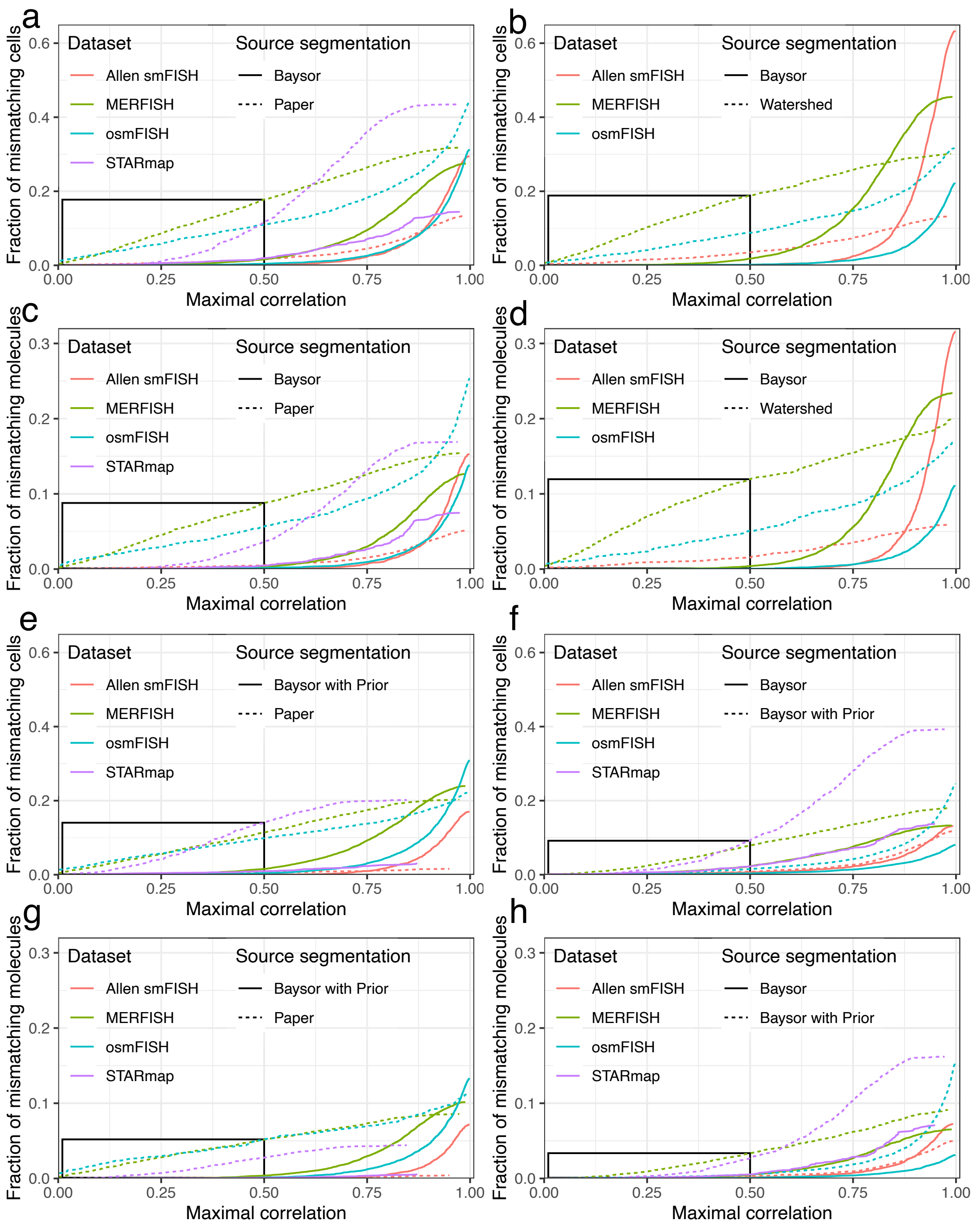


**Supplementary Fig. 9 | Effect size in the correlation benchmarks.** The panel shows the fraction of mismatching cells (**a,b,e,f**) and molecules (**c,d,g,h**) - those for which low correlation between cell parts was observed (see Fig. 4e-f). Specifically, the y-axis shows the fraction of cells/molecules with the correlation between parts below the threshold (x-axis). The fraction is calculated relative to the total number of cells/molecules in the dataset. The line colour denotes the dataset, and the line type denotes the source segmentation. Different pairs of source/target segmentations were evaluated: Baysor against Paper (**a,c**), Baysor against Watershed (**b,d**), Baysor with Prior against Paper (**e,g**), Baysor against Baysor with Prior (**f,h**). The black box in the bottom-left highlights the region with correlation below 0.5, which is the most relevant for benchmarking purposes. It can be seen that incorporating Paper-based Prior brings Baysor segmentation closer to the Paper (**a** *vs.* **e** and **c** *vs.* **g**). On the other hand, incorporating Prior results in higher fraction of mismatching cells and molecules (**f,h**). For the plots **e-f**, the values of the prior segmentation confidence were picked based on the quality of the prior segmentation: Allen smFISH = 0.75, MERFISH = 0.5, STARmap = 0.7, osmFISH = 0.35.

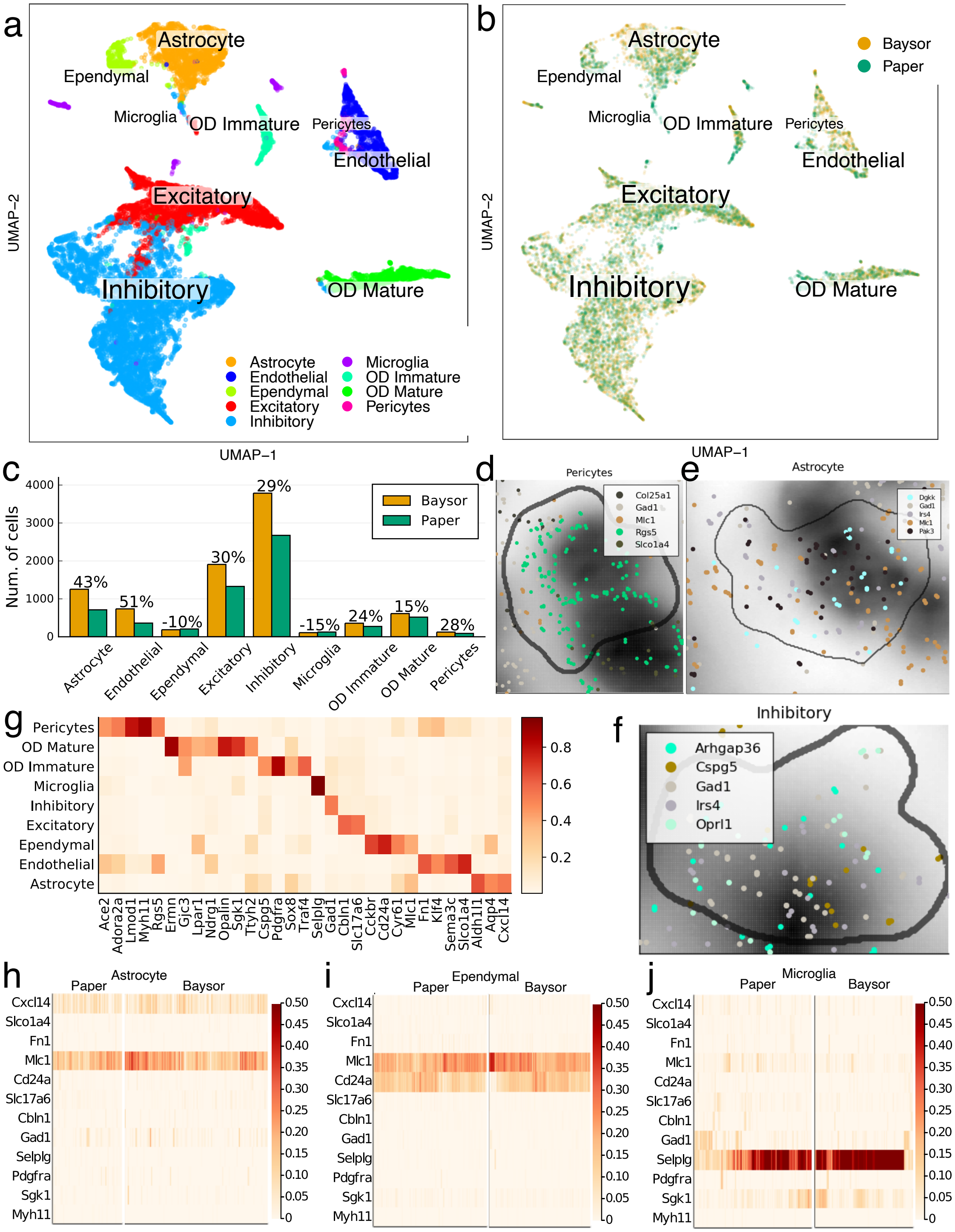


**Supplementary Fig. 10 | Comparison of the Baysor and the published segmentation on the MERFISH Hippocampus dataset.** The figure shows the comparison in the same format as Fig. 5. **a**. A joint UMAP embedding of the cells from both Baysor and the paper segmentations. The colors correspond to the annotated cell types. **b**. The same embedding, colored by the segmentation that produced a specific cell. **c**. The frequency of different cell types is shown for the Baysor (brown bars) and the Paper (green bars) segmentations. The numbers on the top of the bars show excess percentage for Baysor. The largest difference is observed for Endothelial cells, where the Paper segmentation has 51% fewer cells compared to Baysor.

**Supplementary Fig. 10 | d-f**. Examples of Pericytes (**d**), Astrocytes (**e**) and Inhibitory neurons (**f**), which were not segmented by the Paper annotation, but were distinguished by Baysor. The dots correspond to the measured molecules, colored by gene (only five the most abundant genes are shown). The grayscale background shows the DAPI signal, and the black contours show the determined cell boundary. **g**. A heatmap showing expression patterns of marker genes (columns) for the different cell types (rows). The colors show expression levels, normalised for each gene. **h-j**. Expression heatmaps of the key markers for Astrocytes (**h**), Ependymal cells (**i**), and Microglia (**j**). It can be seen that for **h** and **i**, only the marker genes expected in these two cell types are expressed in both Baysor and Paper segmentations. However as can be seen in **j**, the Paper segmentation has higher fraction of cells expressing markers of both Microglia and Inhibitory neurons. These doublets might explain the lower number of cells for the Baysor segmentation.

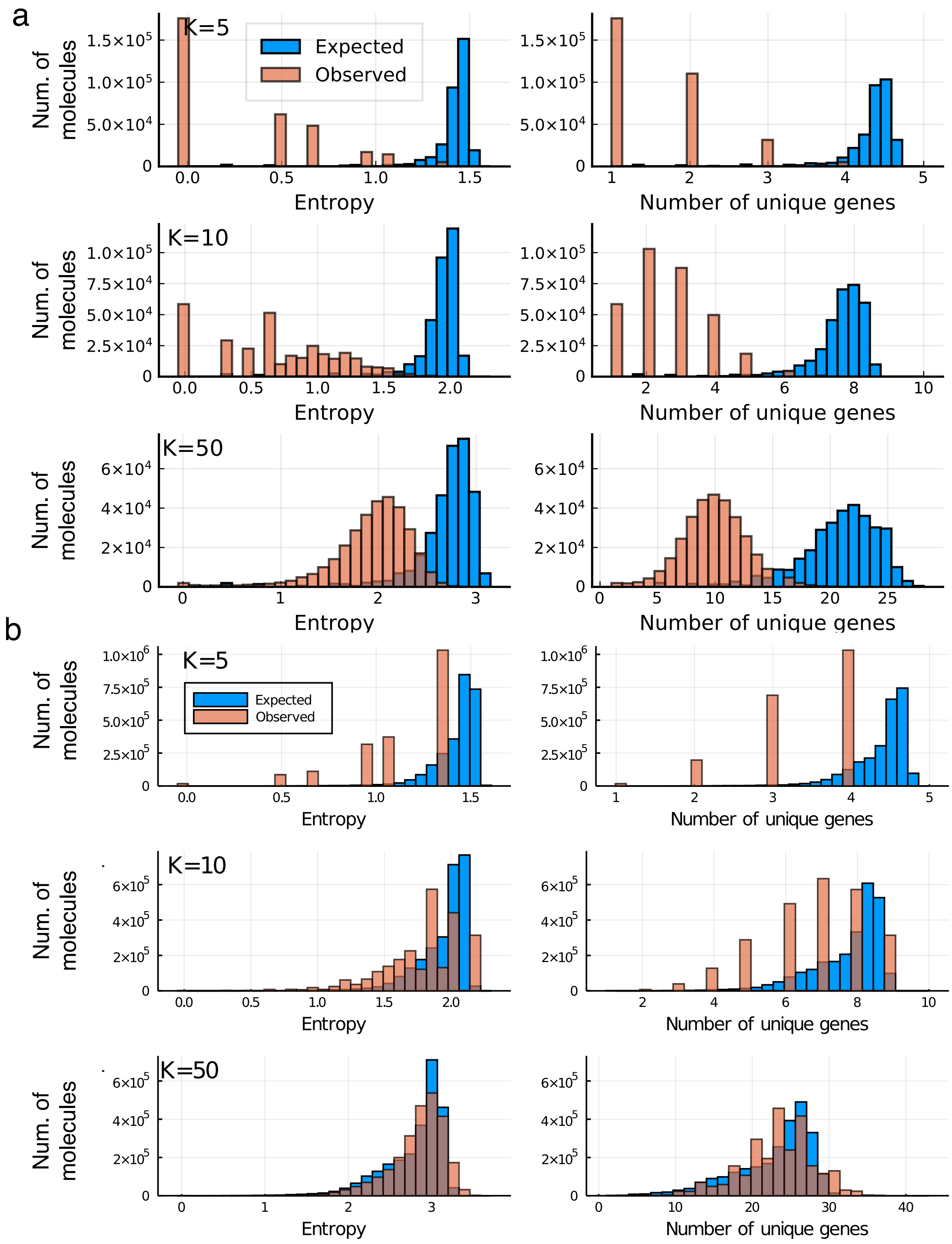


**Supplementary Fig. 11 | Local clustering of molecules on STARmap data.** As in the Fig. 6e of the main manuscript, the figure shows the effect of local grouping on STARmap data (**a**) for different values of $K$ (top-left corner label). The results show lack of a local clustering effect for the MERFISH dataset (**b**) with the expected distribution matching to the observed one.

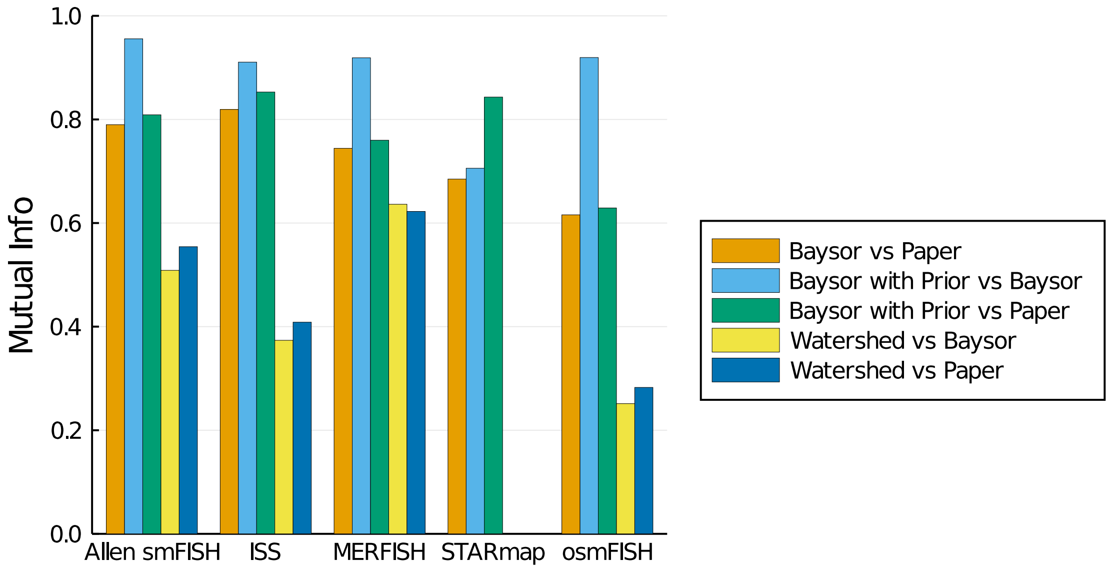


**Supplementary Fig. 12 | Correspondence of different segmentation methods.** Cell assignments from different segmentation methods were compared, and Mutual Information (y-axis) was estimated between the pairs of segmentation results (shown with colors) for different protocols (x-axis). It can be seen that "Baysor with Prior" results are more similar to the published segmentations for all datasets, with the largest difference observed for the STARmap dataset, where Baysor (without prior) performed relatively poorly.

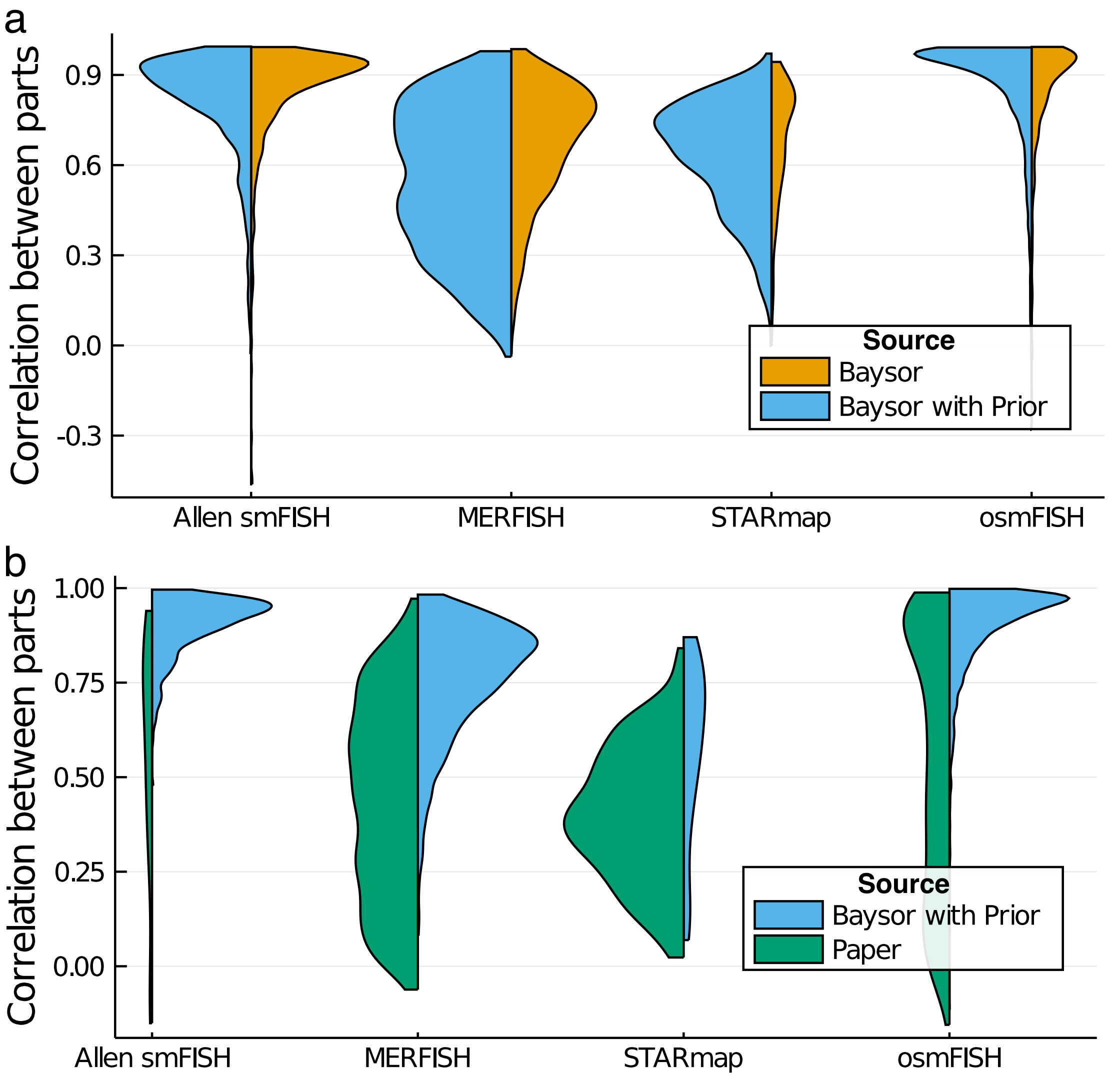


**Supplementary Fig. 13 | Correlation benchmarks for Baysor segmentation with prior.** The correlation benchmarks (see Fig. 4c-f) are shown for the Baysor segmentations without prior against Baysor with Prior (**a**), and for Baysor with prior against Paper (**b**). The format corresponds to the Fig. 4e-f: the correlation distribution is shown on the y-axis, with different protocols arranged on the x-axis. The width of the violin plot is proportional to the number of source cells that were matched to multiple target cells. The benchmark **a** illustrates that "Baysor without prior" shows higher correlation values compared to Baysor with prior. However, it does not necessarily mean that prior should not be used: a staining-based segmentation can resolve situations, which are not captured by the expression correlation benchmark (see "Outstanding challenges" section). The values of the "prior segmentation confidence" were chosen based on the quality of the prior segmentation: Allen smFISH = 0.75, MERFISH = 0.5, STARmap = 0.7, osmFISH = 0.35.

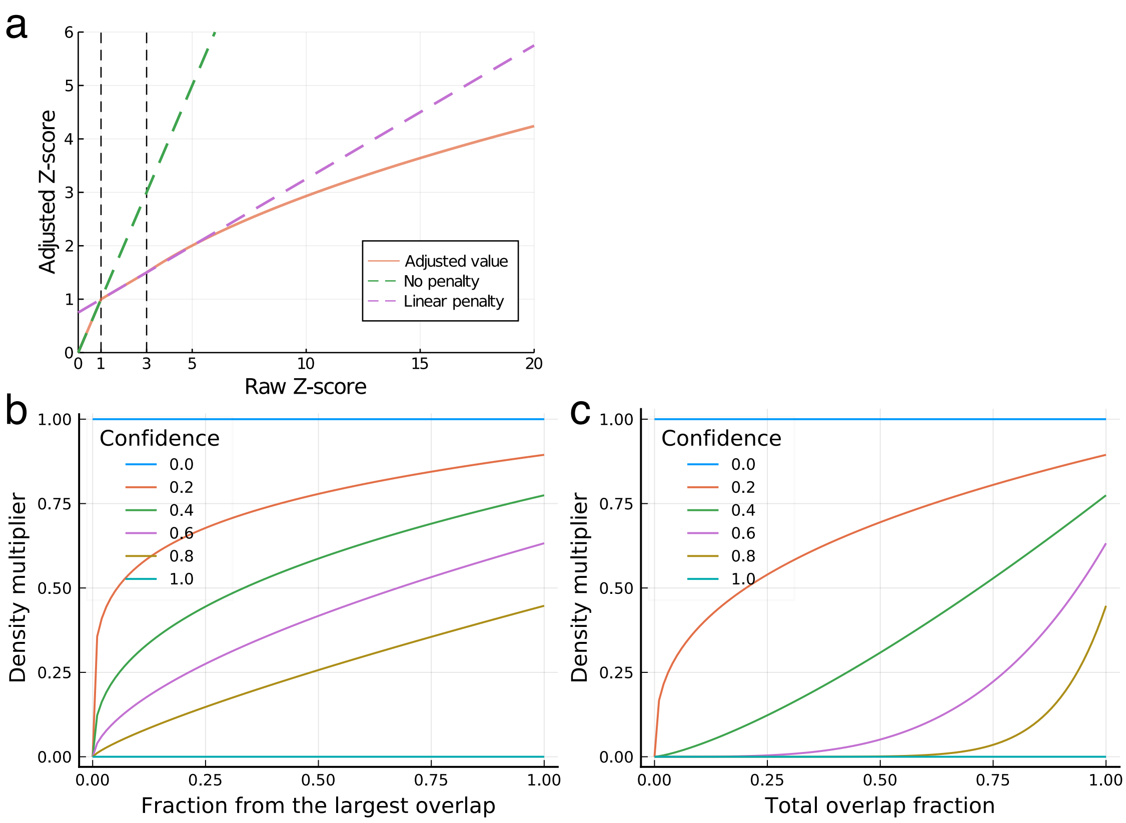


**Supplementary Fig. 14 | Penalty curves for the segmentation algorithm. a**. When transferring annotations from scRNA-seq to spatial molecules, cluster centres are penalised for deviating from the scRNA-seq data in their expression patterns. The penalty depends on the Z-score of the deviation, and the resulting adjustment is structured as follows: no penalty for $Z < 1$; linear penalty for $Z \in [1, 3]$; and super-linear penalty for $Z > 3$. The adjusted Z-score is shown on the y-axis for the proposed penalty function (red), linear penalty (purple dashed line) and no penalty (green dashed line). The black vertical dashed lines show the points where the function changes its shape ($Z = 1$ and $Z = 3$). **b,c**. The penalty for disagreement with a prior segmentation is comprised of two parts: the penalty for two cells being present in the same prior segment (**b**) and the penalty one cell touching multiple prior segments (**c**). The plots show the penalty density multiplier on y-axis as a function of the unwanted overlap fraction ($\frac{n^{seg}_{k,q}}{n^{seg}_{u^*,q}}$ or $\frac{n^{seg}_{k,q}}{n^{seg}_{q}}$ for **b** and **c** correspondingly, see Methods) on x-axis for different prior segmentation confidence values (shown with colors). Both types of penalties start with no penalty for prior confidence of 0.0, and progress to complete density elimination for all but the main component per segment when the prior confidence approaches 1.0 (see Methods for more details on the penalties).

### Supplementary Tables

**Supplementary Table 1. Performance profiling of molecule clustering.** The table shows total runtime for Baysor Markov Random Field (MRF) cell type segmentation and scRNA-seq-like Leiden clustering of Neighborhood Composition Vectors ("CPU time", mean ± standard deviation). MRF-based segmentation is shown for different number of clusters ("Num. clusters"), while for Leiden, the number of clusters does not affect performance. It can be seen that MRF is almost twice as fast as Leiden for 10 clusters, and 5.5 times faster for 2-4 clusters.

**Supplementary Table 2. Performance profiling of Baysor segmentation.** The table shows time and memory profiling for Baysor segmentation on different datasets ("CPU time" and "Max RSS" correspondingly, mean ± standard deviation). It can be seen that the runtime mostly depends on the total number of molecules, while the maximal memory usage is largely determined by the total number of genes.

**Supplementary Table 3. Parameters for Baysor segmentation runs.** The table shows command line parameters for Baysor, used to produce the results for both runs with paper prior (Prior=Yes) and without it (Prior=No). If the parameter was not specified, "NA" value is shown in the table.
